# PlantaeViz: a comprehensive platform for the analysis, integration, and visualization of public omics data across plant and related species

**DOI:** 10.1101/2024.12.19.629382

**Authors:** Antonio Santiago, Luis Orduña, José David Fernández, Álvaro Vidal, Iñigo de Martín-Agirre, Chen Zhang, Purificación Lisón, David Posé, Carmen Martín-Pizarro, Ioannis Kandylas, Iban Eduardo, Elena Monte, M. Águila Ruiz-Sola, Aurélien Devillars, Justin Joseph, Alessandro Vannozzi, Elena A. Vidal, David Navarro-Payá, José Tomás Matus

## Abstract

The increasing availability of omics data from non-model plant species has created a pressing need for centralized, user-friendly platforms that maximize the utility of these datasets in a FAIR manner. Here, we introduce PlantaeViz, a web-based tool designed for the integration, visualization, and analysis of multi-omics data across a wide range of plant species. PlantaeViz offers advanced functionalities, including gene catalogues built from curated literature, transcriptomic meta-analyses presented as gene expression atlases, gene co-expression and regulatory networks with on-the-fly ontology analyses, cistrome visualization and metabolomics-transcriptomics integration, among other tools, providing a robust framework for hypothesis generation and biological interpretation. *Gene Cards* applications, tailored to each plant species, provide both community-curated and automatically generated functional annotation information. One of the platform’s core features is its big data approach: over 58,000 publicly available SRA transcriptomic samples have been processed and visualized to date. Significant efforts have been made in orthology assessment using multiple layers of evidence, as well as in the automatic classification and standardization of omics metadata through regular expressions and data mining. As a result, around 90% of transcriptomic runs have been successfully classified according to sample tissue. These data have been used to construct gene networks via computationally intensive methods based on diverse algorithms. We present a study case to illustrate the platform’s integration and exploratory capabilities. PlantaeViz bridges genomics and functional knowledge between model and non-model plant species and aims to expand its species catalogue of species in the future, democratizing access to large-scale plant omics data. Further developments will include the incorporation of additional data types, and the implementation of new tools to further support plant research across diverse biological contexts.

## Introduction

The rapid accumulation of omics data in non-model plant species has reached levels comparable to those of well-studied model organisms in the recent past. For example, Illumina-based transcriptomic data in grapevine has exceeded 10,000 runs in the NCBI Sequence Read Archive (SRA) (Sayers et al., 2022), a milestone that the Arabidopsis community only reached in 2017. This growth calls for up-to-date, centralized web-based platforms dedicated to these emerging systems. Such platforms are essential for enabling the efficient reuse, integration, and analysis of public datasets. In their absence, the full potential of the knowledge generated by the research community remains underexploited.

While the volume of omics data in non-model crop species now rivals that of model organisms from just a few years ago, the depth of biological insight remains significantly limited. This disparity is largely due to the smaller and more fragmented research communities involved in these species. *In silico* approaches, such as sequence-based functional annotation or gene co-expression analyses, offer powerful strategies to bridge this gap by enabling the prediction of gene functions. Gene co-expression networks, which operate on the principle of ‘guilt-by-association’, have been widely applied in non-model crops (Childs et al., 2011; Liesecke et al., 2019; Orduña et al., 2023). In these networks, genes that are consistently co-expressed across diverse conditions are more likely to share functional relationships, even if their specific roles remain unknown. As such, co-expression networks provide a valuable framework for inferring the functions of poorly annotated genes based on their association with better-characterized counterparts.

The integration of multi-omic datasets, spanning genomics, transcriptomics, and metabolomics, is central to systems biology, offering a holistic understanding of organismal function. Yet, despite their potential, these datasets are often underexploited due to the absence of accessible, intuitive tools for their visualization, exploration, and reanalysis. Publicly available omics data can reveal complex gene expression patterns across tissues, developmental stages, and environmental conditions, providing insights into regulatory mechanisms and phenotypic traits that frequently transcend the original scope of the study. Unlocking this latent value requires platforms that empower both experts and non-specialists to mine and reinterpret existing datasets in meaningful ways.

Model organisms such as *Arabidopsis thaliana* benefit from comprehensive, well-maintained databases, notably The Arabidopsis Information Resource (TAIR) (Huala et al., 2001). TAIR integrates omics resources through standardized gene nomenclature, high-quality genome assemblies, and curated structural and functional annotations. It also offers a suite of tools for exploring large-scale datasets, including access to mutant collections, literature-based gene function curation, and gene expression profiles. In contrast, existing databases for non-model plants are often limited in scope and functionality. They frequently lack advanced features such as gene co-expression network analysis or tissue- and condition-specific expression exploration. These are essential tools for effective hypothesis generation and systems-level research. Moreover, broad cross-species platforms like STRING (Szklarczyk et al., 2023), while valuable, tend to underperform for less-characterized plant species, as they rely on outdated annotations, legacy gene identifiers from obsolete older genome assemblies or annotations, or microarray probe-specific identifiers, which limits their utility for more modern, RNA-seq-based studies.

In this study, we present PlantaeViz (https://plantaeviz.tomsbiolab.com/), a versatile and centralized platform for the visualization and analysis of functional, and genomic data across a range of plant species. The platform is organized into modular suites, which currently include grapevine (*Vitis vinifera*), tomato (*Solanum lycopersicum*), mulberry (*Morus alba*), *Cannabis sativa,* and *Arabidopsis thaliana* in its initial release. Unlike model organism-centric resources, PlantaeViz incorporates the latest genome assemblies and annotations for these species. Its development has involved extensive manual curation, including literature-based compilation of gene catalogues and the creation of hierarchical ontologies to support consistent metadata classification across datasets. Notably, the platform integrates *Arabidopsis* data as a functional gateway, employing its well-annotated genome to facilitate comparative insights into less-characterized species. Future releases are planned to expand the platform with additional species suites, in response to evolving research priorities. PlantaeViz features an intuitive, user-friendly interface equipped with built-in tools to generate heatmaps, line plots, and co-expression network diagrams, designed to support data interpretation and hypothesis generation. By addressing the lack of dedicated and integrative resources for non-model plants, this platform empowers plant researchers to maximize the utility of omics datasets and accelerate functional gene discovery in key agricultural and medicinal crops.

## Results

### A modular framework for expanding species and tools in plant omics

PlantaeViz (https://plantaeviz.tomsbiolab.com/) consolidates a broad and diverse set of transcriptomic resources across several plant species of agronomic and medicinal relevance: grapevine, tomato, mulberry, and cannabis. Arabidopsis is included as a reference, to allow for easier transfer of functional knowledge from model to non-model species. The platform is organized into different suites that can be accessed from the homepage; *Vitis spp.* (VitViz), *Solanum lycopersicum* (TomViz), *Morus spp.* (MulViz), *Cannabis spp.* (CannaViz), and *Arabidopsis thaliana* (AraViz). The suites follow a modular structure designed for continuous expansion, facilitating the adaptation of the platform for additional species, several of which are currently under development, including *Chlamydomonas reinhardtii* (ChlamyViz), *Prunus spp.* (PrunViz), *Fragaria spp.* (StrawViz), among others.

Each suite offers a homepage that features an informative index with menus linking to various applications (apps) and tools, including gene network visualization, gene expression atlases, differential gene expression analysis, gene set enrichment analysis, genome browsers, multiple-sequence alignment, gene function annotation, and an updated BLAST server (Figure 1). Interoperability between the different suites has been enabled through comprehensive orthology analyses between species, cultivars and/or genome versions. These multi-tool analyses include a summary output and orthologs have been assigned high, medium or lower confidence based on a custom scale.

**Figure 1.**
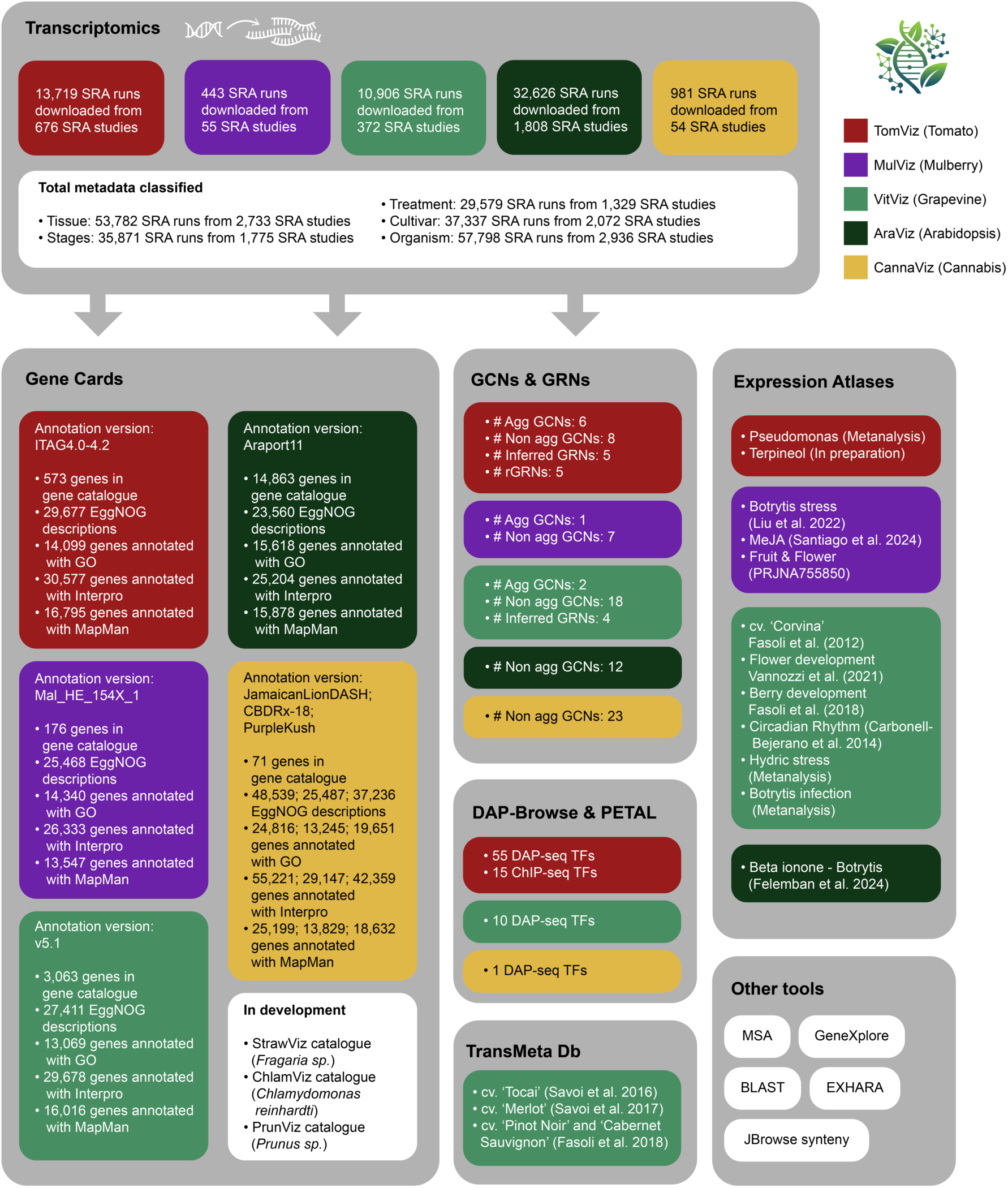
Overview of the content and functionalities in the different suites of the PlantaeViz platform. Publicly available, reannotated transcriptomic datasets are used by the *Gene Cards* module to generate tissue-specific expression profiles for each species, and by the Gene Co-expression Networks (GCNs) and Gene Regulatory Networks (GRNs) modules. PlantæViz also includes: *DAP-Browse* and *PETAL* (Plant Expression and Transcription factor Association Lookup) apps, for exploring genome-wide DNA-binding peaks; *TransMetaDb*, for integrated transcriptome-metabolome analyses; Expression Atlases providing curated, study-specific and meta-analysis overviews; and a collection of auxiliary tools for sequence alignment, functional annotation and comparative genomics.

The platform integrates a vast collection of Illumina-based transcriptomic data, over 57,000 SRA runs, which have been systematically categorized by metadata categories such as tissue type, developmental stage, treatment condition, cultivar, and organism. A custom set of hierarchical ontologies, coupled to regular expressions, has been designed to achieve this. These transcriptomic datasets are the basis for multiple functional modules within PlantaeViz, enabling both gene-level exploration and systems-level analyses.

### Automated classification of transcriptomic data using custom species ontologies

Automatic classification of SRA runs was performed for five key metadata categories (cultivar, organism, stage, tissue, and treatment type), across five analyzed species (grapevine, tomato, mulberry, cannabis, and arabidopsis). This classification was based on species-specific ontology frameworks developed for each organism (Supplemental Data 1-10, in both JSON and pdf format). The resulting classifications are presented in Supplemental Tables 1-5, displaying all terms automatically identified by the pipeline (in the “*_curated_details*” columns) alongside the single, most specific assigned term (in the “*_curated*” columns). In cases where terms from distinct ontology branches (e.g., ‘fruit’ and ‘root’), were simultaneously detected, a single-term assignment was not feasible and such entries were labeled as “mix”. Assignment confidence for each term was also provided on a scale of 0-10.

Overall, assignment success varied considerably across categories (Figure 2A). The “tissue” category showed one of the highest success rates, with nearly complete annotation coverage in most species. This high performance can be attributed not only to the common practice of submitting tissue-related metadata, even when not standardized, but also to the robustness of our ontology-driven approach. The tissue ontologies were carefully adapted for each species, accounting for anatomical and terminological differences, which contributed to more accurate classification results.

**Figure 2.**
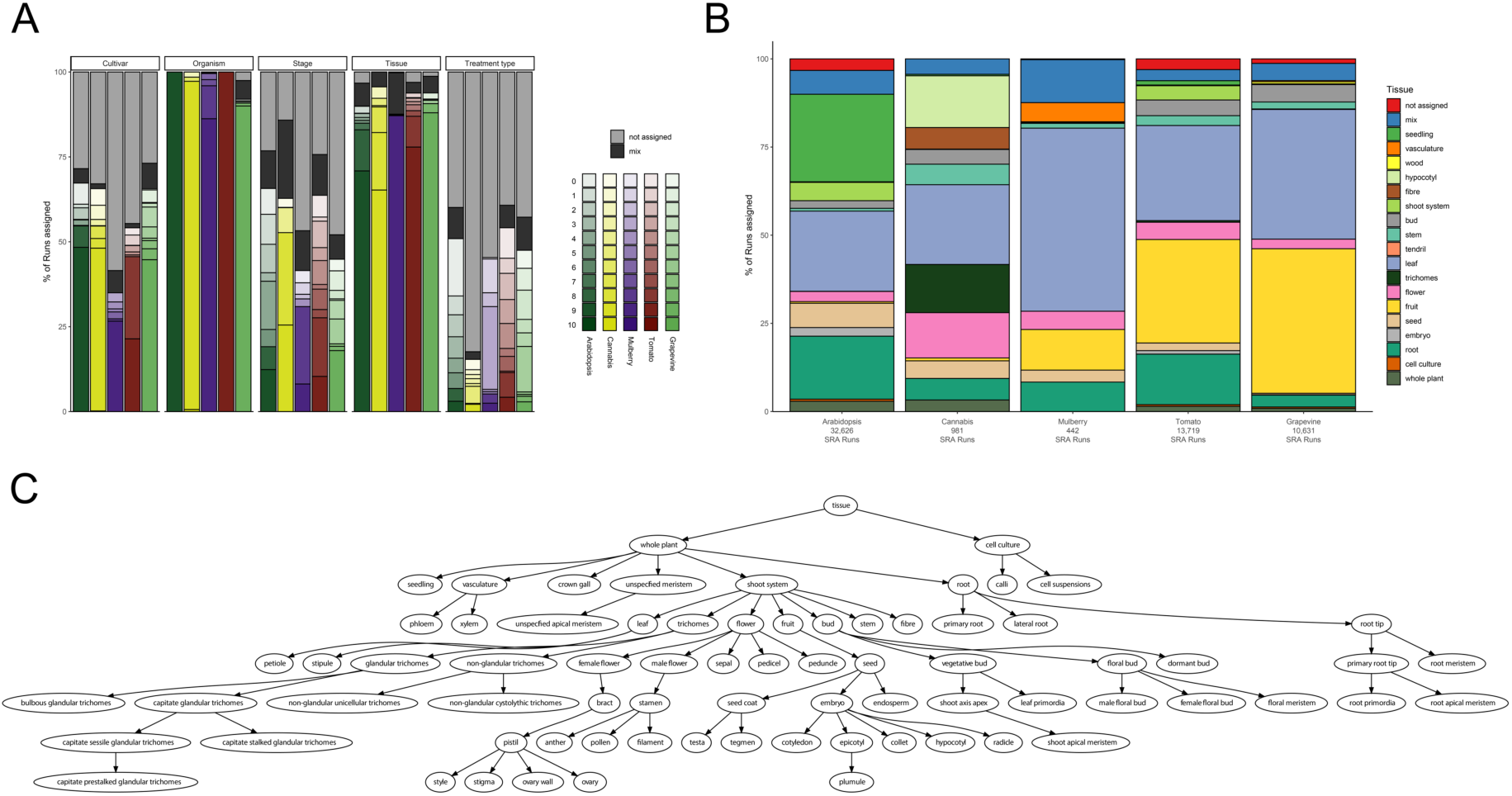
Ontology-driven reannotation of SRA runs across metadata categories and plant species. (A) Percentage of SRA runs successfully assigned to metadata categories across different species, alongside their corresponding confidence level on a 0-10 scale. Lighter tones indicate assignments made with lower confidence according to a custom scale based on a combination of metadata column and regular expression reliability. The *tissue* and *organism* categories show the highest assignment success across all species. (B) The distribution of tissue-type assignments varied substantially between species, reflecting differences in tissue representation that align with species-specific research priorities. (C) Example of the tissue ontology for *Cannabis sativa*, illustrating hierarchical relationships between terms. This structure enables flexible grouping of SRA runs into broader categories, even when the sampled tissue corresponds to a specific subcomponent. Full graphical representations of all ontologies can be found in Supplemental Data 6-10.

Analysis of the specific tissue types assigned revealed distinct patterns reflecting the predominant research focus for each species (Figure 2B). In the model organism *A. thaliana*, the most frequently represented tissues were leaf, root, and seedling, with notable contributions from vascular tissues and cell cultures, indicative of its broad experimental use. In cannabis, flower and trichome tissues dominated, consistent with a research focus on specialized metabolite biosynthesis. For grapevine, tomato, and mulberry, the majority of samples were classified as leaf or fruit, aligning with studies on fruit development and metabolic regulation. Notably, root tissues were markedly underrepresented in grapevine, comprising only 3.4% of runs, compared to 14.3% in tomato. This discrepancy may stem from the fact that *Vitis vinifera* is commonly studied for its aerial organs, while rootstocks used in viticulture often belong to other *Vitis* species, which are less frequently included in transcriptomic analyses.

Similar to the “tissue” category, the “organism” category achieved near-complete assignment success, reaffirming that species-level information is generally well represented in public repositories. In contrast, assignment rates for the “cultivar” and “stage” categories were more variable (Figure 2A). For instance, mulberry cultivars appear to be more sparsely annotated as the percentage of assigned runs falls below 40% whilst the other species surpass 50%. Similarly, developmental stage information was assigned to over 50% of runs in tomato, cannabis and arabidopsis, but remained lower for grapevine and mulberry, potentially reflecting greater heterogeneity or incompleteness in the available metadata. The “treatment type” category, which is still under refinement, reached approximately 50% assignment success in some species. This is encouraging, particularly given that not all transcriptomic studies involve treatments, and associated metadata is often inconsistently reported.

These results underscore the inherent challenges in curating and standardizing unstructured metadata, particularly for non-model species where such information is often poorly annotated. While our ontology-based approach (Figure 2C) significantly improved the reannotation of SRA runs, its performance was ultimately contingent on the quality and completeness of the metadata available for each species. Figure 2C illustrates the hierarchical structure of tissue terms in *Cannabis sativa*, reflecting species-specific anatomical and developmental distinctions, such as separate categories for male and female flowers, and a refined classification of globular trichomes based on their reported developmental stages and morphological forms (Livingston et al., 2020). As a further example, cannabis fruits are technically achenes; however, since the use of “fruit” was more frequently used in the original metadata and both terms are correct, we adopted “fruit” as the primary ontology term, with regular expressions configured to recognize both “achene” and “fruit” as valid synonyms.

In total, we downloaded and processed 32,626 transcriptomic samples for *A. thaliana*, 13,719 for *S. lycopersicum*, 10,631 for *V. vinifera*, 981 for *C. sativa*, and 442 for *M. alba* (Supplemental Tables 1-5). Expression data from samples with successfully assigned tissue annotations were integrated into the *Gene Cards* and *GeneXplore* applications, enabling intuitive visualization and exploration. A curated subset of these annotated datasets was further used in downstream analyses, including co-expression network generation or regulatory network inference.

### A unified resource for gene-centric insights

The *Gene Cards* application provides gene-centric information across all species, incorporating automatic functional annotation for queried genes, structural models integrated via *JBrowse2*, ortholog inference, TPM-based expression profiles across all metadata-annotated RNA-seq tissue samples from SRA, and access to gene, transcript, CDS, and protein sequences (Figure 3).

**Figure 3.**
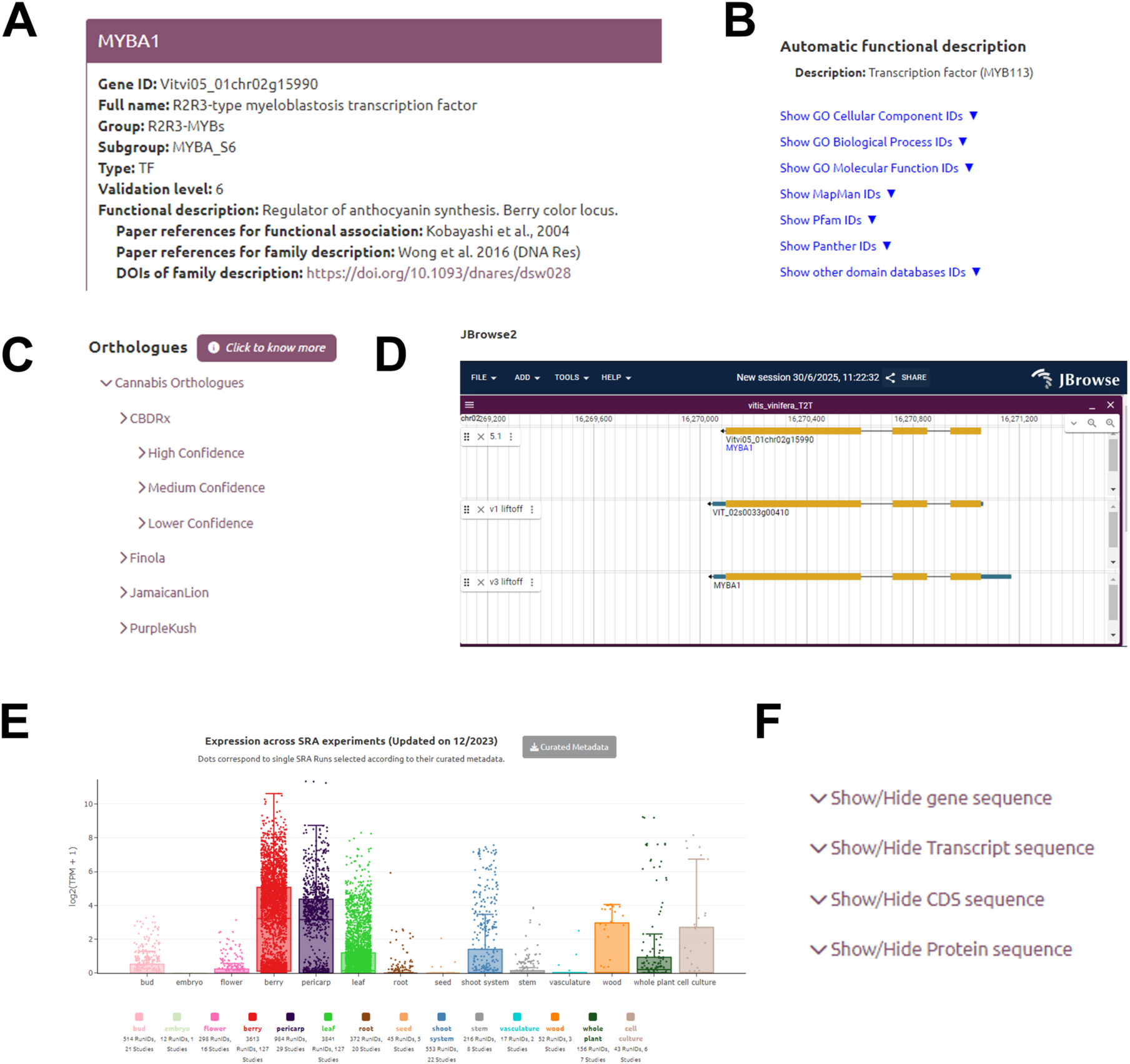
The grapevine gene *VviMYBA1* as an example in VitViz *Gene Cards* app. (A) Manually curated grapevine gene catalogue information. (B) Automated functional annotation with structured, category-specific dropdown menus. (C) Predicted orthologs across other species and cultivars included in PlantaeViz split by confidence level. (D) Embedded *JBrowse2* instance displaying the genomic context of the selected gene. (E) Tissue-specific expression profile of the gene across more than 10,000 SRA runs, shown in log₂(TPM+1). The interactive boxplots allows users to identify the corresponding SRA studies by hovering over individual data points. (F) FASTA-format sequences for the gene, transcript, coding sequence (CDS), or protein are also available.

Each *Gene Cards* app functions as the main access point to a curated gene catalogue, allowing users to retrieve detailed and curated information for individual genes. Grapevine is currently the best-curated species within the platform, and similar curation efforts are underway for other plant species to broaden the platform’s utility. In the case of this species, it is tightly linked to the grapevine gene reference catalogue (Navarro-Payá et al., 2022) (available at https://grapedia.org/genes/). This curated resource, currently in its third version, includes functional data for over 3,000 genes and is open to community submissions through the Grapedia portal. The gene catalogue is built upon the foundation of the telomere-to-telomere (T2T) v5 cv. “PN40024” genome assembly, where v5.1 gene IDs serve as the primary identifiers for genes. Provided information includes gene symbols and synonyms, descriptive gene names, pathway and gene family classifications, and custom-defined validation levels for functional data. To facilitate the research community’s transition to updated genome annotations, the platform includes a cross-annotation tool that enables users to input one or more gene IDs and retrieve the corresponding identifiers across all supported annotation versions, ensuring backward compatibility and smooth interoperability.

### Interactive visualization of transcriptomic data

The platform incorporates expression atlas applications, which are built upon manually curated datasets specifically designed to address distinct research needs. These applications consolidate transcriptomic data from various studies, encompassing both microarray and RNA-seq datasets and allowing to investigate gene expression patterns across a wide array of biological contexts, including specific tissues, developmental stages, and experimental treatments. Examples include atlases for specific flower developmental stages (Vannozzi et al., 2021), hormone elicitation (Santiago et al., 2024) or the circadian rhythm (Carbonell-Bejerano et al., 2014), as well as different meta-analysis atlases combining multiple experiments regarding treatments, such as hydric stress, and *Botrytis* infection (Felemban et al., 2024; Liu et al., 2022). Among their key features, users can generate interactive heatmaps based on custom gene input lists, apply data transformations such as logarithmic or Z-score scaling, and perform clustering to group data by genes or conditions, thereby highlighting co-expression patterns. Expression atlases support the visualization of normalized transcript levels (e.g. TPM, transcripts per kilobase of exon model per million mapped reads) (Figure 4A) as well as differential expression data (Figure 4B). In differential expression views, heatmap cell color represents fold-change values, while dot size conveys adjusted p-values, facilitating intuitive interpretation of significance and magnitude. These features collectively enable robust comparative analyses, allowing researchers to detect expression trends and infer potential functional associations under varying biological conditions.

**Figure 4.**
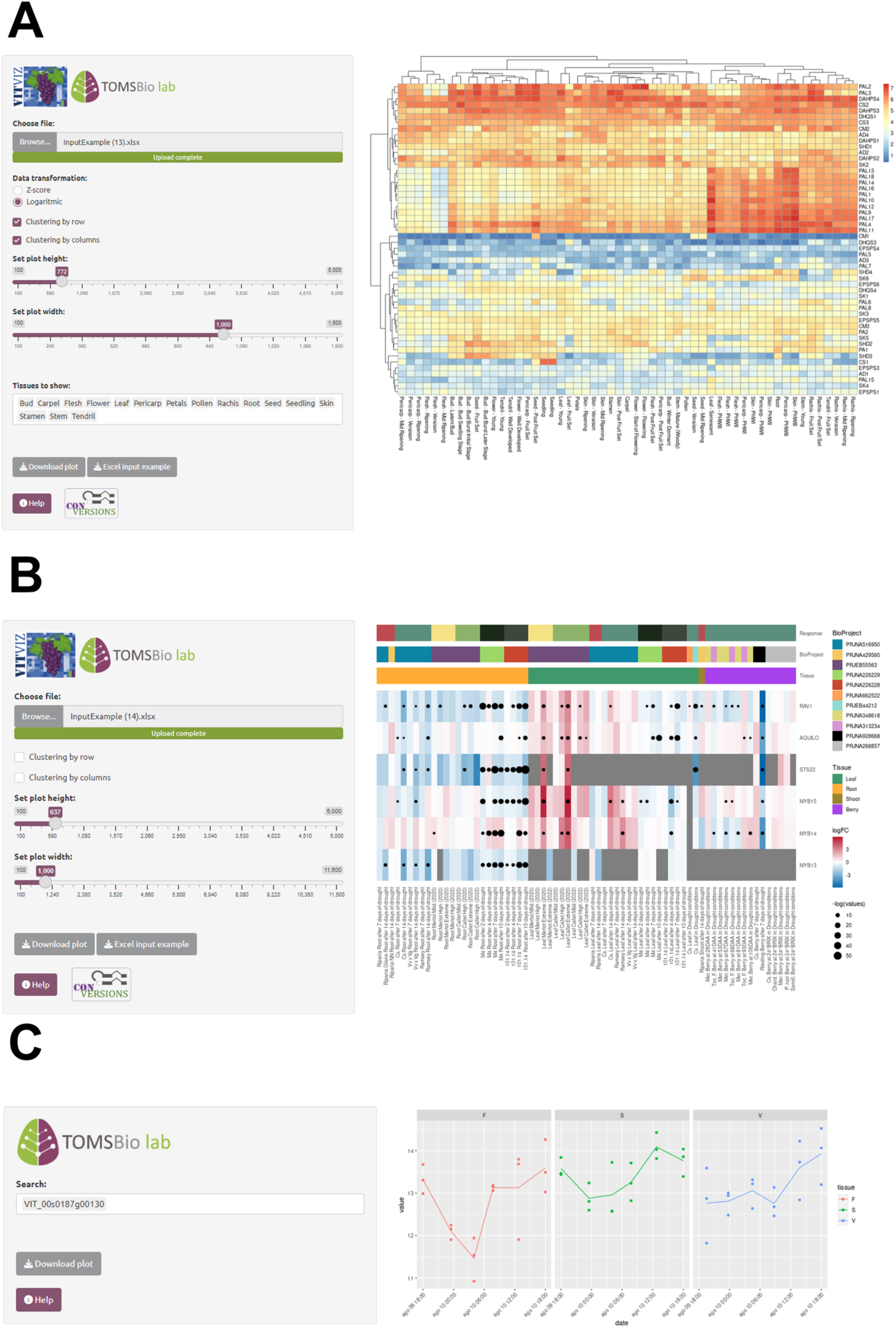
Examples of gene expression atlases integrated into PlantaeViz. (A) A detailed view of the *Vitis vinifera* cv. ‘Corvina’ expression atlas based on normalized microarray data, showcasing gene expression patterns across 54 developmental stages and tissue types (Fasoli et al., 2012). (B) Differential gene expression results compiled from various independent transcriptomic studies focused on hydric stress conditions. This meta-analysis allows users to explore gene-specific responses to drought and water deficit treatments across genotypes, organs, and experimental setups. (C) Normalized microarray gene expression at different timepoints for grapevine cv. ‘Tempranillo’ flesh (F) and skin (S).

For further data exploration, we developed two complementary web applications: *EXHARA* (EXpression Heatmaps Across all SRA Runs Available) and *GeneXplore*. *EXHARA* is designed for the broad visualization of gene expression from a user-provided list of genes across all curated runs (Supplemental Figure 1A). Its primary output is an interactive heatmap that can be dynamically filtered by tissue or by specific user-selected runs, providing a versatile resource for customized analysis. Alternatively, *GeneXplore* was developed to integrate automatic metadata classification results with gene expression data in a user-friendly web application that allows visualization and comparison of gene expression levels across a custom range of experimental categories (Supplemental Figure 1B). The tool integrates the ontology-based annotations described above and displays normalized TPM expression values enabling cross-sample comparability. *GeneXplore* allows users to examine expression differences for specific genes across tissues, developmental stages, and treatment conditions, individually or in a combinatorial manner, based on the curated and available metadata. Together, these tools provide a powerful resource for uncovering patterns and gaining insights into gene behavior across diverse biological contexts.

### Exploring gene-to-gene relationships through network-based analyses

PlantaeViz hosts a comprehensive collection of over 90 whole-genome networks, including aggregated and non-aggregated gene co-expression networks (GCNs), as well as gene regulatory networks (GRNs), inferred either through co-expression or supported by experimental evidence. Among these, GCNs are particularly amenable to AUROC-based performance evaluation, as the ‘guilt-by-association’ principle underpins their structure and predictive utility (Supplemental Table 6). In this study, we applied the GCN construction methods previously developed by Orduña et al. (2023), ensuring methodological consistency with prior benchmarking efforts. Analysis reveals that network performance improves with increasing number of studies and/or sequencing runs, regardless of species or GCN construction method (Supplemental Figure 2A). This trend supports the benefit of dataset breadth and diversity in co-expression network reliability. Notably, aggregated GCNs almost always outperform non-aggregated networks, showing marked improvements in predictive power across species (Supplemental Figure 2B).

We have developed interactive visualization tools for exploring gene co-expression networks (GCNs) (Figure 5A) and gene regulatory networks (GRNs) (Figure 5B), each combining multiple functionalities to investigate gene relationships and regulatory interactions. The applications are organized into two main tabs, each tailored to facilitate different aspects of network analysis while presenting advanced visualization features for in-depth exploration. By combining co-expression data with regulatory predictions, including transcription factor (TF) binding events at open chromatin regions, the platform enables construction and interrogation of regulatory networks. This dual approach enables users to retrieve TF-target gene pairs, visualize specific subnetworks based on custom gene sets, and explore co-expression and regulatory interactions concurrently. Notably, the underlying network data are not static; they are periodically updated to incorporate new experimental datasets.

**Figure 5.**
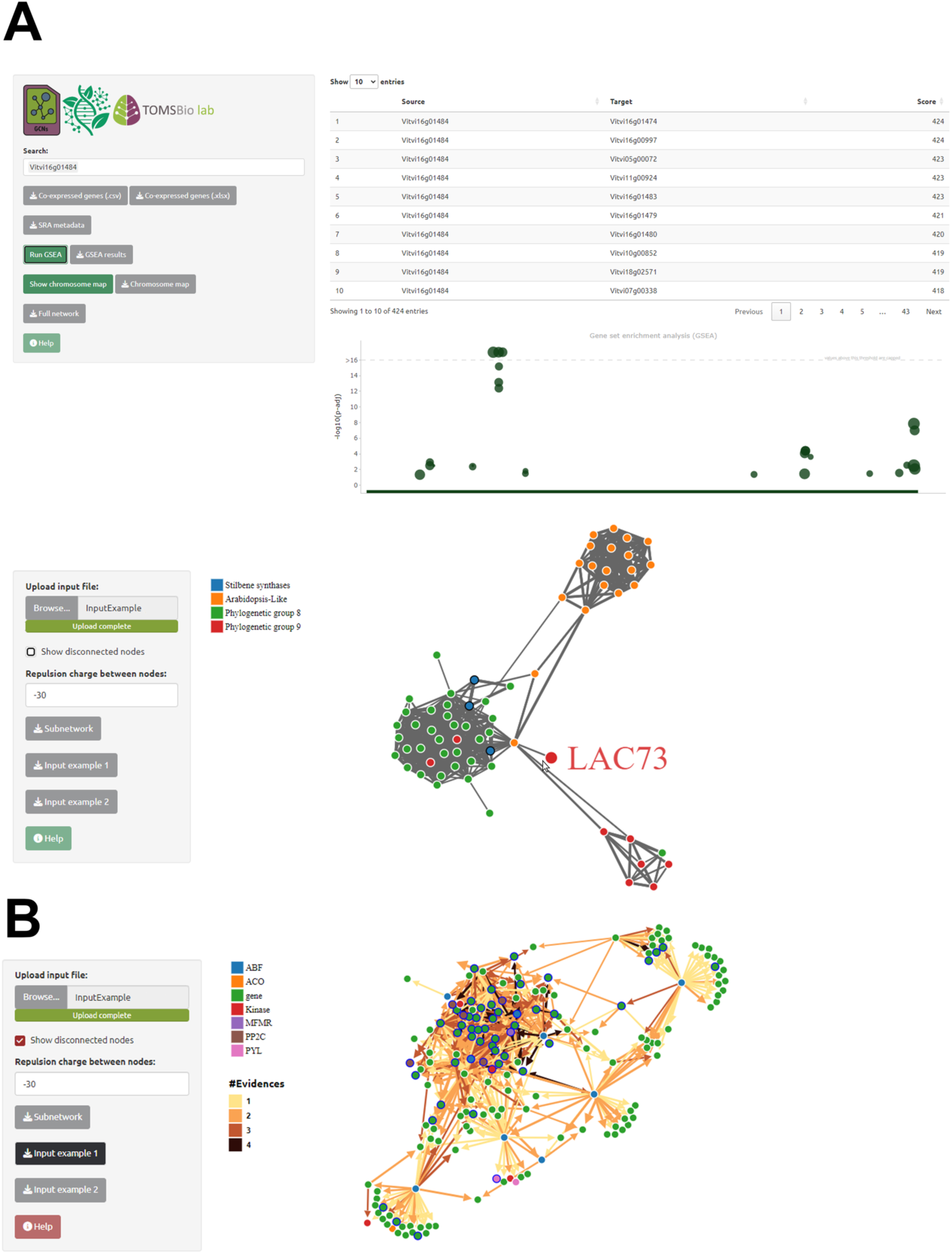
Examples of gene co-expression network (GCN) and gene regulatory network (GRN) applications in PlantaeViz. (A) The GCN app provides a list of genes co-expressed with a selected gene of interest based on large-scale transcriptomic data. Users can input a gene identifier to retrieve co-expression partners and visualize their relationships using an interactive network plot. This plot highlights the strength and direction of co-expression correlations, allowing users to identify potential functional associations or gene modules. Additionally, users can submit a custom list of gene IDs to generate interaction plots, facilitating the visualization of known or hypothesized networks. The figure displays the AggGCN App interface within the VitViz module of PlantaeViz. (B) The GRN app enables the visualization of predicted regulatory interactions between transcription factors (TFs) and their target genes. The example shows an interaction plot where edges represent putative regulatory relationships inferred from expression data and/or experimental binding evidence. Nodes are color-coded by gene function or regulatory role, providing an intuitive overview of gene network topology and regulatory hubs. The figure displays the GRN App interface within the TomViz module of PlantæViz.

The application interface includes a co-expression table with chromosomal mapping, as well as a 2D force-directed network visualization module. In the first tab of GCN apps, users can query a gene of interest to retrieve a list of co-expressed genes that is by default filtered to the top 1% of interactions, this is otherwise known as a gene-centered network. This filter can be optionally skipped to display all co-expression relationships present in the whole network, to better reflect the scale-free topology of biological networks. While most nodes are expected to have few connections, a few nodes will be highly interconnected, i.e. not all nodes will have the same number of edges within the whole network. This table provides detailed information such as gene identifiers and names (sourced from curated gene catalogues), along with co-expression metrics. Users can perform on-the-fly gene set enrichment analysis (GSEA) on the retrieved list to identify enriched biological pathways or functional categories. Additionally, the chromosomal locations of the co-expressed genes are displayed in an integrated genomic map, enabling the exploration of their genomic distribution. This feature facilitates the identification of potential chromosomal clustering or co-localization of functionally related genes, bridging expression-level analysis with genomic context. GRN apps offer similar functionality, however, gene search can be focused on source, target genes, or both. This reflects the fact that source genes in a GRN are always TFs and target genes may be TFs or otherwise, thereby allowing to query potential regulators as well as regulatory targets.

The second tab offers an interactive 2D force-directed graph visualization of gene co-expression or regulatory networks based on the network D3 R package. A key feature is the ability to input custom gene lists, enabling users to explore specific gene sets and their interconnections based on co-expression patterns or regulatory interactions. The network is rendered as a dynamic, two-dimensional graph in which nodes represent genes and edges denote co-expression or regulatory relationships. This visualization is especially valuable for exploring pathway- or family-oriented gene networks, helping to uncover potential co-regulation and connectivity among genes of interest. Users can identify hub genes (i.e., potential key regulators), and detect tightly connected clusters within their input set. The topology of the D3 subnetwork is influenced by node connectivity: node positions are a compromise of push/pull forces, they repel each other by default and attract each other based on number edges and their weights. The layout is also interactive, allowing the user to identify gene names and edge weights by simply hovering over. Moreover, some visual rearrangement of nodes is allowed, after which forces are recalculated, always maintaining a force-directed topology. Users can adjust repulsion parameters to optimize layout visualization according to the characteristics of their subnetwork. To ensure analytical reproducibility in light of these periodic updates, each downloaded metadata includes a timestamp, allowing users to reference the specific network version used for their visualization and analysis. In compliance with FAIR principles, the full network can be downloaded at any time as well as a comprehensive set of metadata that describe each of the transcriptomic studies used to build it and any relevant network building parameters. NCBI data download timestamps and transcriptomic query dates are provided for traceability, accounting for the fact that further aggregation of SRA runs to these networks will occur in future updates, thereby changing the networks topology.

### Exploratory applications and genome browsers for ChIP-seq and DAP-seq data

Understanding transcriptional regulation is essential for dissecting traits such as development, stress response, and fruit quality. Chromatin immunoprecipitation (ChIP)-seq and DNA-affinity purification (DAP)-seq resources facilitate the exploration of transcription factor (TF)-DNA interactions across plant species and represent a very good, yet partially complete, view of the the regulatory landscape. This data is presented in the form of JBrowse adaptations, here named *ChIP-/DAP-Browse*s, which integrate community-contributed ChIP- and DAP-seq datasets, acting as an interactive platform for visualizing cistromes and DNA-binding profiles (Figure 6A). A summary of currently available transcription factors is provided in Supplemental Table 7, including both our own and publicly available data. The collection is expected to grow as further contributions are added by the research community.

**Figure 6.**
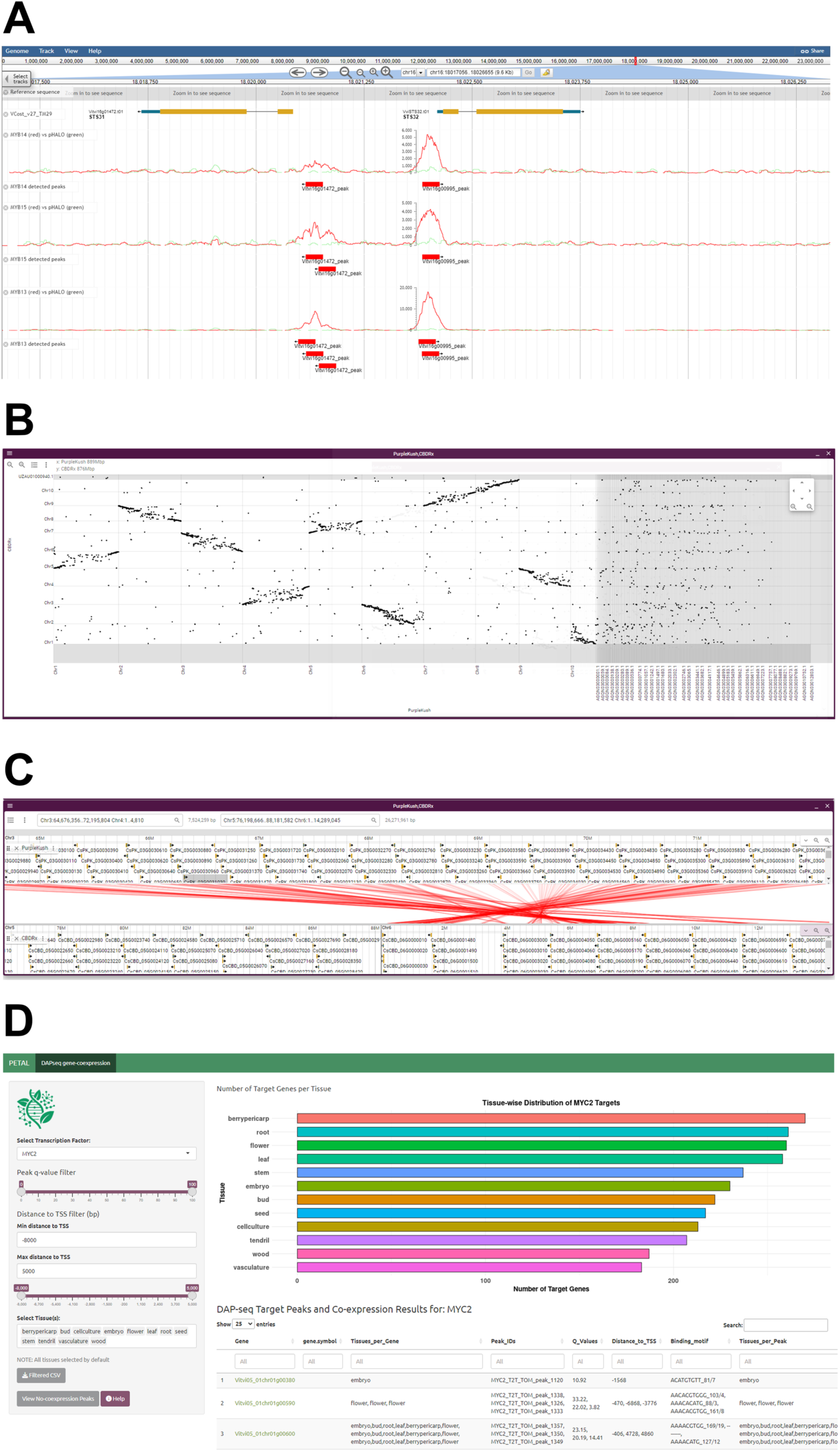
Examples of *ChIP/DAP Browse* and gene- and protein-centered module of *PETAL* app. (A) Genome-wide DNA-binding data displayed in the *DAP-Browse* module, showing DAP-seq peaks associated with stilbene synthase genes targeted by the transcription factors *VviMYB13, VviMYB14 and VviMYB15* in *Vitis vinifera*. The viewer allows inspection of binding site locations relative to gene features, enabling hypothesis generation on transcriptional regulation. (B) Dotplot comparison between the *Cannabis sativa* cultivars cv. ‘CBDRx’ (y-axis) and cv. ‘Purple Kush’ (x-axis), illustrating sequence similarity and large-scale structural variation across the two genomes. (C) Detailed synteny view between a region in chromosome 06 of CBDRx and its homologous region in chromosome 03 of Purple Kush. Gene models within these regions are linked based on conserved collinearity, providing insight into genome evolution and gene conservation across cultivars. (D) The protein-centric tool within *PETAL* integrates DAP-seq binding data with tissue-dependent and independent GCN information, MYC2 is shown as an example query. The interface displays a dynamic bar plot summarizing the tissue-wide distribution of co-expressed gene targets for a selected TF, based on user-defined filters such as peak quality (q-value) and distance to TSS. A responsive table provides detailed target information, including gene identifiers (hyperlinked to catalogue information within the *Gene Cards* apps), ChIP/DAP-seq peak metrics, and a list of tissues where TF–target co-expression is detected. Together, these components enable tissue-specific prioritization of regulatory interactions.

Genome annotation tracks follow a standardized color scheme: coding exons are represented as yellow boxes, untranslated regions (UTRs) as blue boxes, and introns as connecting lines. For each transcription factor, *ChIP-/DAP-Browse* provides three types of visualization tracks: 1) Individual replicate tracks, showing raw alignment distributions for both TF and input libraries, 2) Merged replicate tracks, allowing comparison of aggregated TF versus input signal, and 3) GEM peak-caller tracks, highlighting predicted binding peaks specific to each TF. Unlike static tabular peak data, which may be prone to false negatives or positives due to algorithmic limitations, the genome browser enables manual inspection of read alignments, allowing users to visually assess peak quality and binding signal consistency. GEM peak-caller tracks also associate detected peaks with their most likely target genes, using a nearest-genomic-feature assignment method. In addition to TF binding data, the genome browser includes dot plot visualizations for inter-genome comparisons (Figure 6B), and interactive synteny maps displaying conserved gene relationships across species (Figure 6C).

In order to complement *JBrowse2* capabilities and to delve deeper into cistrome data, we also introduce *PETAL* (Plant Expression and Transcription factor Association Lookup), an interactive app that allows users to explore DAP-seq data from both gene-centric and protein-centric perspectives.

The gene-centered tool (Supplemental Figure 3) allows to examine DAP-seq peaks from multiple TFs associated with multiple genes in a single view. Peaks are represented within a user-defined distance from the transcription start site (TSS) of their corresponding genes. Users can also optionally filter peaks based on fold-change thresholds. The shape of each peak is represented by plotting the centers of reads from the bigWig files (x-axis) against their scores (y-axis). To standardize the display, maxima and minima are set equal, and other points are scaled accordingly. Due to this, absolute peak heights are not comparable between different peaks, but fold-change, derived from the annotation, is conveyed through peak color intensity.

On the other hand, the protein-centered tool (Figure 6D) integrates DAP-seq data with the previously presented GCNs, to draw gene regulatory networks and identify putative targets for TFs of interest. This combined dataset, derived from both ChIP/DAP-seq binding profiles and co-expression patterns, allows users to prioritize tissue-specific regulatory links by filtering targets based on TF binding peak quality, proximity to transcription start site (TSS), and tissue expression. The interface provides a drop-down menu of TFs, accompanied by sliders and numeric filters to adjust q-value thresholds and maximum distance to TSS, enabling users to focus on high-quality peaks and gene targets relevant to the TF’s specific role. Results are visualized as a dynamic bar plot summarizing the number of co-expressed genes per tissue type. The bar plot updates interactively as the user adjusts filters, providing insight into the reliability and tissue specificity of the associations. Detailed results are displayed in a dynamic, filter-responsive table listing gene targets with their IDs, gene symbols (where available), ChIP/DAP-seq peak information (including q-value and distance to TSS), and co-expressed tissues. Gene IDs in the table are hyperlinked to detailed gene information within the gene catalogue and the *Gene Cards* app. For gene targets that are also co-expressed with the TF, the table lists the tissues in which co-expression is observed, providing a comprehensive view of regulatory interactions. All of the above tools offer a robust and user-friendly framework for investigating transcription factor binding dynamics and their regulatory roles across diverse plant genomes.

### Integration of gene-metabolite data across experiments

*TransMetaDb* is an interactive tool designed for the integrated visualization and analysis of transcriptomic and metabolomic datasets (Supplemental Figure 4). It enables users to explore correlations between gene expression levels and metabolite abundances across diverse experimental conditions. By facilitating the identification of functional associations, *TransMetaDb* helps uncover potential gene-to-metabolite relationships that may underlie specific physiological or biochemical traits. The primary output is a downloadable heatmap or table, where rows represent user-input genes and columns correspond to different metabolites. Each cell reports the Pearson’s correlation coefficient (PCC) along with the associated p-value (shown in brackets), calculated using the Student’s asymptotic method. This matrix allows rapid visual and statistical assessment of correlation strength and significance. Additionally, the tool features a module for exploring gene co-expression clusters derived from weighted gene co-expression network analysis (WGCNA), performed locally. Users can download a complete table assigning all genes to their respective modules, enabling further exploration of gene-metabolite associations within co-regulated gene sets. This dual functionality provides a powerful framework for systems-level analysis of metabolic and transcriptional correlation.

### Other more common tools: multiple sequence alignment

To facilitate the exploration of sequence conservation and functional relationships among homologous genes, we included a multiple sequence alignment (MSA) application, enabling users to align nucleotide (mRNA or CDS) or protein sequences, helping identify conserved domains, motifs, and sequence variations that provide insights into gene function and evolutionary dynamics (Supplemental Figure 5). The application supports a list of gene IDs for the species of interest and offers multiple alignment algorithms, including ClustalW, Clustal Omega, and MUSCLE. These supported methods allow customization of the alignment process to meet specific analytical needs. Alignment results are displayed in an interactive, color-coded viewer. Conserved residues or nucleotides are highlighted, and a corresponding sequence logo graphic summarizes conservation patterns across the alignment. Users can zoom in on specific regions for detailed inspection and export results in FASTA format for downstream analyses. By integrating flexible alignment options with intuitive visualization, the MSA application offers a streamlined solution for studying sequence conservation, functional domains, and genetic variation across related gene sets.

### Case study: Interspecies functional association of MYB24 in terpene biosynthesis

This case study demonstrates the utility of the GCNs applications, cistrome browser, and ortholog inference tools available in the platform to perform integrative cross-species regulatory analyses. Specifically, we investigated the functional conservation of the MYB24 transcription factor in regulating terpene biosynthesis in *Vitis vinifera* and *Cannabis sativa*.

We first used DAP-seq data to identify direct TF-DNA interactions. In *V. vinifera*, the DAP-seq assay for *VviMYB24* (Zhang et al., 2023) revealed 3,557 high-confidence binding events located within 5 kb upstream of transcription start sites (TSSs), while *CsMYB24G* showed 1,926 such binding events (Zhang et al., unpublished results). Consistent with their potential regulatory roles in reproductive tissues, both genes showed high expression levels in flower samples, as displayed in the *Gene Cards* application (Figure 7A).

**Figure 7.**
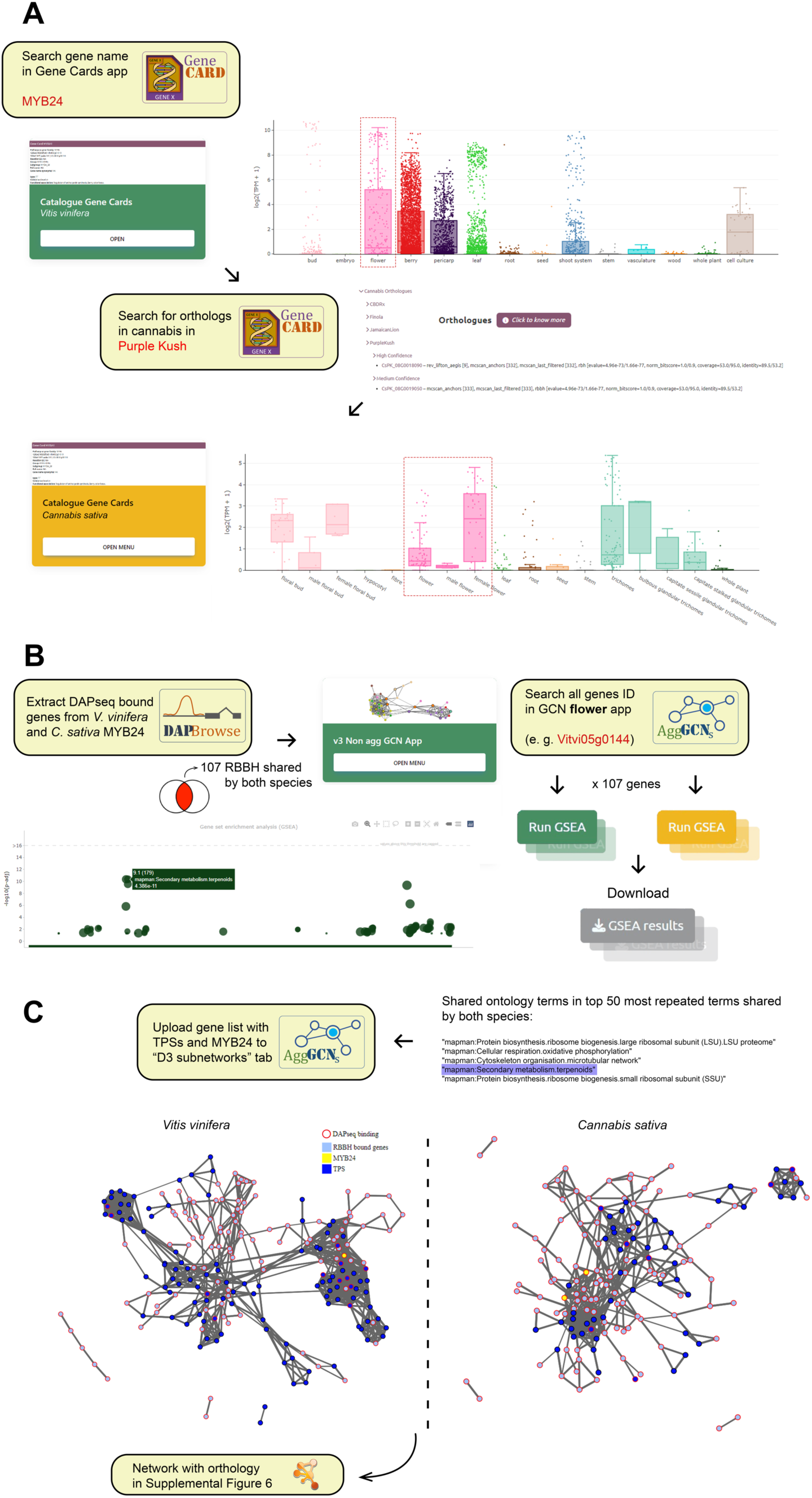
Cross-species validation of MYB24 regulatory function using gene coexpression networks (GCNs) and DAP-seq data using PlantaeViz. (A) Tissue-specific expression profiles of *MYB24* in *Vitis vinifera* and its ortholog *MYB24G* in *Cannabis sativa* based on publicly available SRA transcriptomes visualized through the *Gene Cards* app, highlight conserved high expression in floral tissues. (B) Gene set enrichment analysis (GSEA) of a DAP-seq-bound gene, conducted using the “Co-expressed Genes” tab in the GCN app. Each dot represents an enriched Mapman term linked to orthologous co-expressed genes (via Reciprocal Best BLAST Hits, RBBH) in both species, illustrating conserved regulatory pathways. (C) Cross-species co-expression subnetworks were generated via the “D3 Subnetworks” tab, using manually curated terpene synthase (*TPS*) genes and DAP-seq targets as inputs. Networks show MYB24 targets and their co-expression relationships with *TPS* genes in *V. vinifera* (left) and *C. sativa* (right). Nodes with red borders indicate direct DAP-seq binding events, supporting a degree of functional conservation across species.

To compare targets across species, we used the platform’s built-in reciprocal best BLAST hit (RBBH) ortholog mapping between *C. sativa* and *V. vinifera*. We identified 107 genes bound by *MYB24*/*MYB24G* with 1-to-1 orthology across the two species. These shared DAP-seq targets were further analyzed for functional context by performing gene set enrichment analysis (GSEA) on their top co-expressed partners, using the “Co-expressed Genes” tab of the GCN app (Figure 7B). To improve specificity and exclude overly generic terms, GO categories with more than 200 annotated genes were filtered out.

Remarkably, in both species, the fourth most frequently enriched term among the co-expressed gene sets, after more general categories like LSU proteasome, oxidative phosphorylation, and microtubule network, was associated with terpene biosynthesis. This was the top-ranked term specific to specialized metabolism, indicating a potentially conserved regulatory function of *MYB24* in controlling terpene production.

To further investigate the network connectivity among *MYB24* targets and terpene biosynthesis genes, we curated a list of terpene synthase (*TPS*) genes from both genomes. These TPSs were combined with the DAP-seq derived targets and submitted to the “D3 Subnetworks” visualization tool in the platform (Figure 7C). This allowed us to reconstruct co-expression subnetworks and inspect the architectural organization of *MYB24*-associated modules highlighting direct and indirect associations between MYB24 targets and *TPS* genes. The graphical outputs revealed both conserved and species-specific co-expression structures. The resulting networks for *V. vinifera* and *C. sativa* were also exported as CSV files and visualized using Cytoscape (Supplemental Figure 6). Together, these findings support the hypothesis of a conserved MYB24-centered regulatory module involved in floral terpene biosynthesis, while also reflecting evolutionary divergence in network topology and regulatory wiring between these two species.

## Discussion

PlantaeViz distinguishes itself from existing omics databases through its unique integration of diverse functionalities, its emphasis on non-model plants, scalability, and a user-centered design. The platform unifies transcriptomic, genomic, and metabolic datasets into a single coherent framework, enabling multi-omics exploration across a growing number of plant species. It has been engineered to handle large-scale datasets efficiently, with improved webpage responsiveness and robust data-handling capabilities. Extensive manual curation and intuitive analytical interfaces allow users to perform species-specific or comparative analyses. Applications such as *Gene Cards* and Expression Atlases present a detailed view of gene expression dynamics, while tools like *TransMetaDb* and interactive gene network visualizations provide insights into functional associations between genes, metabolites, and regulatory elements.

Compared to existing platforms, PlantaeViz offers a more integrative, up-to-date, and FAIR-compliant approach to plant genomics and transcriptomics. Many widely used resources remain static, siloed, or limited in scope and accessibility. For example, TAIR, while historically invaluable for Arabidopsis research, has transitioned to a restricted-access model, limiting open data availability; something fully addressed in the open-access AraViz suite of PlantaeViz (Huala et al., 2001; Reiser et al., 2024). Similarly, Sol Genomics and TomExpress provide rich resources for Solanaceae (Fernandez-Pozo et al., 2015; Zouine et al., 2017), but TomViz offers a curated re-annotated gene models and improved catalogues that are regularly updated for accuracy and completeness. VitExpress, in contrast, is built on a non-reference grapevine genome and lacks curated gene catalogues or gene network tools (Djari et al., 2024), whereas VitViz utilizes the latest T2T reference genome and integrates co-expression and regulatory networks using ChiP- and DAP-seq data. Even large-scale platforms such as Phytozome, despite their broad range of genomes available, do not support transcriptomic analyses or functional network visualizations (Goodstein et al., 2012). PlantaeViz surpasses these limitations by offering a dynamic, integrative environment that unifies up-to-date genomic and transcriptomic data with advanced analytical tools, making it not just a complement but a necessary next-generation platform for plant systems biology.

Another key distinguishing feature of PlantaeViz is its extensive transcriptomic data coverage across multiple plant species. In its initial release, the platform integrates a total of 58,351 accessions from the NCBI SRA database, which has been instrumental in developing advanced tools such as expression profiling and regulatory network visualizations. This achievement was made possible through the use of high-performance computational resources, ensuring that the platform remains efficient and scalable as dataset complexity grows. The processed transcriptomic data has been used to build tissue-specific co-expression networks for all included species, and high-performing aggregated gene co-expression networks (AggGCNs) were generated wherever data volume permitted. While network aggregation has been demonstrated to improve network performance, it requires substantial datasets and computational power (Wong, 2020).

A further critical advantage of PlantaeViz lies in its modular and species-agnostic design. This flexible architecture ensures broad compatibility with a wide range of plant species and facilitates the integration of additional organisms and emerging computational methods over time. By supporting both well-established and non-model plants, interoperability across knowledge bases is enhanced, improving data accessibility and analytical capabilities for underrepresented species while maintaining essential links to well-characterized model organisms. The platform’s capacity to integrate multi-omic datasets, including transcriptomics and metabolomics, further highlights its value for functional genomics and hypothesis generation. Its suite of analytical tools enables the exploration of functional relationships, identification of conserved gene regulatory motifs, and analysis of gene-metabolite correlations, providing researchers with a systems-level understanding of plant biology. This is particularly advantageous for advancing studies in plant development, specialized metabolism, and environmental responses, where interest in non-model species is rapidly growing.

Looking ahead, the ongoing incorporation of updated datasets, additional species, and new analytical methods will continue to expand PlantaeViz’s utility. By embracing the latest multi-omics technologies and broadening its species coverage, the platform is poised to remain a central resource for systems biology research, facilitating novel discoveries and accelerating translational applications in agriculture and biotechnology.

## Materials and methods

### Web server implementation

PlantaeViz was developed as a Shiny server (v1.5.17.973) running R version 3.6.3. The index page is based on React as the JavaScript library. The platform is freely available for use on the web and can be found at https://plantaeviz.tomsbiolab.com/.

### Download of public transcriptomic data and metadata classification

Raw RNA-Seq data were retrieved from the Sequence Read Archive (SRA), using the SRA Toolkit with a specific focus on Illumina-based sequencing runs. Only transcriptomic datasets were included, excluding those related to microRNAs, small RNAs, or non-coding RNAs. Species-specific queries were designed to identify relevant RNA-Seq experiments, with the full list provided in Supplemental Table 8. In cases where species-level annotations were limited, genus-level queries were used to capture data from closely related accessions. For *Arabidopsis thaliana,* an additional filter was applied to include only paired-end sequencing studies given the high volume of available data.

Associated metadata was automatically classified using a custom species-specific ontology. This ontology, stored as a structured JSON file, includes hierarchical terms across five main categories: tissue, organism (species), developmental stage, cultivar and treatment type. Each term is linked to a curated set of search terms and regular expressions used to detect their presence within relevant metadata fields (Supplemental Table 9). Only columns relevant to a given category and species were considered during classification. To assess classification confidence, a qualitative scoring scale ranging from 0 to 10 was implemented (Supplemental Table 10). Scores were assigned based on two criteria: (1) the reliability of the metadata field (e.g., dedicated column vs. abstract) and (2) the specificity of the regular expression used (categorized as ‘reliable’ or not). A score of 9 represents high-confidence assignments, where a reliable regular expression matches a dedicated reliable metadata field. In contrast, a score of 0 indicates a low-confidence match, typically from general text fields using less specific expressions. A score of 10 was reserved for manually curated annotations.

### Development of *Gene Cards* applications

Functional annotation was generated for each species using both eggNOG-mapper and Mercator4, extracting GO and MapMan terms, Pfam, Panther and other databases domains, and then reformatting GO and MapMan ontologies to GMT format for gprofiler2 GSEA analysis. In the case of grapevine, curated gene functional information was integrated based on the submission forms provided by the research community (https://grapedia.org/genes/). Gene ID correspondences between grapevine annotations were established by using the annotation extraction and gene integration suite (AEGIS), which was developed *in-house* and is based on a weighted multi-feature coordinate overlap algorithm. A *JBrowse2* genome browser was set up for each species considering the latest genome assembly and annotation versions. An integrated *JBrowse2* window was included using the inbuilt language for gene coordinate queries.

### Construction of gene co-expression (GCN), inferred regulatory (iGRN) and multi-layered regulatory (rGRN) networks

The network construction involved a series of steps to collect, process, and analyze RNA-Seq data obtained from the NCBI SRA database, which was independent of the type of network built; non-aggregated gene co-expression network (non-agg GCN), aggregated GCN (agg GCN), or gene regulatory networks (GRNs). All RNA-Seq reads were processed in a series of steps to prepare them for alignment to the reference genome. This involved trimming the reads using the fastp (version 0.20.0) (default parameters + *--cut_front_window_size 1 --cut_front_mean_quality 30 --cut_front cut_tail_window_size 1 cut_tail_mean_quality 30 --cut_tail -l 20*) to remove low-quality sequences and adapter contamination. Subsequently, the trimmed reads were aligned against the corresponding genome assembly for each species (Table 1) using the STAR aligner (version 2.7.3a) (default parameters + *--runMode alignReads -- limitOutSJcollapsed 8000000 --limitIObufferSize 220000000*). Following alignment, raw count matrices were generated for each SRA run using featureCounts (version 2.0.0) and the respective annotation version for the species (Table 1). To account for variations in sequencing depth across different runs, the raw count matrices were normalized using the transcripts per kilobase of exon model per million mapped reads (TPM) method. Genes exhibiting consistently low expression levels, defined as less than 0.5 TPM across all runs within a study, were removed from further analysis.

**Table 1.** Criteria for ortholog confidence tier classification. Combinations of evidence required to classify an orthologous gene pair into one of three confidence tiers: High, Medium, or Lower. These tiers integrate multiple lines of evidence, including coordinate-based lift-over (AEGIS score), sequence homology (RBBH, RBH), and synteny (MCScan) to provide a final reliability assessment for each identified ortholog pair. RBBH: Reciprocal Best BLASTp Hit; RBH: Filtered Reciprocal BLASTp Hit.

| Confidence Score | Conditions to Achieve this Reliability Level |
| --- | --- |
| High confidence | An AEGIS (Liftoff/LiftOn) score of 7 or greater must be present. |
| Medium confidence | Any of the following conditions are met: (RBBH), (AEGIS score $\geq 6$ and RBH), (RBH and MCScan synteny). In essence, this level requires strong evidence like RBBH, or a combination of slightly weaker evidence types (RBH, MCScan, and moderately high AEGIS scores). |
| Lower confidence | Any of the following conditions are met: RBH is present (but not combined with other required evidence for a higher tier), MCScan synteny is present (but not combined with other required evidence for a higher tier), Other evidences like Orthofinder or lower AEGIS scores exists but does not meet any of the criteria for medium or high confidence. |

The process of constructing aggregated (agg) and non-aggGCNs involved a combination of correlation analysis and network aggregation techniques as described in (Orduña et al., 2023) with some modifications in the pipeline that permit SRA study tracking for aggGCNs. In the case of aggGCNs, the measure of co-expression is based on the number of SRA studies where a co-expression relationship can be ascertained for any pair of genes. Knowing the nature of these studies and not only the total number can be particularly useful to uncover potential condition-specific relationships. Network performance evaluation was carried out for all GCNs using the EGAD R package, whose resulting metric is an AUROC (area under the receiver operator curve) (Ballouz et al., 2017).

The construction of inferred GRNs (iGRNs), were based on the GENIE3 algorithm implemented in the GENIE3 R package (Huynh-Thu et al., 2010). Reference GRNs (rGRNs) included additional *in silico* and experimental sources of evidence to qualify regulatory relationships into 4 confidence levels. The D3 network R package was modified and implemented within the different network apps to show interactive 3D plots based on user-defined gene lists, allowing for focused exploration of co-expression relationships within a specific set of genes. A custom slider to modify repulsion forces is provided. The chromosome map utility was custom built with ggplot2. D3 subnetwork visualization for GRNs was modified to allow for directional edges in the form of arrows (suggestive of regulation) and a color-coded system was implemented for rGRNs to indicate the different levels of evidence supporting a particular relationship.

### Implementation of a genome browser with DAP-seq data: *DAP-Browse*

Sequencing reads obtained from DAP-seq and ChIP-seq experiments were aligned to the corresponding species reference genome using Bowtie2 (Langmead & Salzberg, 2012) with default settings. Reads with MAPQ scores below 30 were filtered out during post-processing to ensure data quality. Peak detection was carried out with the GEM peak caller (Guo et al., 2012) using the respective genome assembly and specifying the following parameters: -q 1 -t 1 -k_min 6 -kmax 20 -k_seqs 600 - k_neg_dinu_shuffle. Analysis was limited to nuclear chromosomes, and replicates were processed using GEM’s multi-replicate mode. Peak summits identified by GEM were mapped to the nearest gene models from a custom annotation file using the BioConductor package ChIPpeakAnno (Zhu et al., 2010) under default parameters (NearestLocation).

To facilitate downstream analysis and visualization, the detected peak coordinates were converted to a BED file format. Alongside the alignment-derived bigWig file, these data were uploaded into the *JBrowse2* genome browser for integration and inspection.

### Utilizing the cross-species synteny visualization features of *JBrowse2*

Pairwise ortholog identification was conducted across all species. For each unique species pair, coding sequences (CDS) were extracted and their corresponding GFF3 annotation files for each species were processed to extract the gene models. Only primary transcripts were retained. Orthologous relationships between species pairs were inferred using MCscan (jcvi.compara.catalog ortholog module) from the JCVI utility libraries and incorporated into the genome browser.

### Differential expression analysis and metabolite integration

Transcriptomic and metabolomic integration was performed as described for the *TransMetaDB* app (Savoi et al., 2022). Briefly, expression data was normalized to TPM. Differential expression analysis (DEA) was performed using the LIMMA R package. Genes were classified as differentially expressed when the adjusted p-value (Benjamini-Hochberg correction) was below 0.05. Co-expression network analysis was performed using the Weighted Gene Correlation Network Analysis (WGCNA) R package for metabolite integration.

### BLAST servers for custom cross-species interoperability

Each species’ datasets were individually formatted into BLAST-compatible databases using the *makeblastdb* utility. We implemented a local installation of SequenceServer (v3.1.3) (Priyam et al., 2019), an intuitive and user-friendly platform for running BLAST (basic local alignment search tool) queries on transcriptomic, CDS, and protein data. The user is able to select parameters based on the BLAST command line tool. Output formats include alignments, detailed hit descriptions, and graphical overviews of query matches.

### Ortholog identification and confidence classification

Orthologous gene relationships were identified using a multi-evidence strategy that integrates synteny, sequence similarity, and annotation lift-over methods to maximize both confidence and genome coverage. For each candidate ortholog pair, evidence from all applied methods was aggregated to support robust classification.

Coordinate-based orthology was inferred using two annotation lift-over pipelines: Liftoff (Shumate & Salzberg, 2021), designed for closely related species or alternative assemblies, and LiftOn (Chao et al., 2024), optimized for more distantly related species, which augments Liftoff with Miniprot (Li, 2023) to guide gene model transfer via protein similarity. In both pipelines, annotations were transferred reciprocally between each genome pair, from source to target genome and *vice versa*. Resulting gene models were then analyzed using our custom tool, AEGIS (annotation extraction genomic integration suite), which detects orthologs by comparing coordinate overlaps between lifted and native gene models. Gene pairs that successfully mapped to each other in both directions were considered reciprocal orthologs, while those matching in only one direction were classified as unidirectional. Each pair was assigned an AEGIS score (1-11) to reflect the quality of the lift-over match, and was annotated with additional flags indicating local synteny conservation (’synteny’) or the matching through duplicate gene models (’copies’) from the Liftoff/LiftOn outputs.

Synteny-based orthologs were identified using the MCscan module from the JCVI toolkit, which detects collinear genomic blocks between species. Two output sets were generated: a high-confidence “anchors” set, and a broader “last_filtered**”** set based on less stringent criteria. Sequence homology was assessed through multiple approaches. First, OrthoFinder (Emms & Kelly, 2019) was used to cluster proteins into orthogroups and infer gene-level orthology using evolutionary gene trees, with the orthogroup identifier serving as supporting evidence. Additionally, all-versus-all BLASTp searches between proteomes were conducted to identify several categories of sequence similarity. Among these, Reciprocal Best BLASTp Hits (RBBH) were defined strictly as protein pairs where each is the other’s top hit, without further filtering. A broader category of Filtered Reciprocal BLASTp Hits (RBH) was also established, retaining reciprocal pairs that satisfied thresholds of E-value ≤ 1e-5, percent identity ≥ 30%, and alignment coverage ≥ 30%. For each BLASTp-based hit, the platform reports additional metrics including E-value, percent identity, percent coverage, and a length-normalized bitscore to support comparative evaluation. Finally, all sources of orthology evidence were integrated and used to assign ortholog pairs into three confidence tiers based on agreement across methods (Table 1).

### Big data processing through high performance computing

Processing the large volumes of transcriptomic data and generating gene co-expression networks required access to high-performance computing (HPC) infrastructure. For this purpose, the Garnatxa HPC cluster at I2SysBio was utilized. The cluster comprises 464 CPU cores and 14 TB of RAM, with peak usage reaching up to 200 CPU cores and 1.2 TB of RAM simultaneously. Efficient handling of these tasks was achieved through the parallelization of Sequence Read Archive (SRA) downloads and data processing pipelines using the SLURM workload manager. This computational capacity was essential to ensure timely and scalable processing of the extensive dataset integrated into PlantaeViz.

## Data availability

In compliance with FAIR (findable, accessible, interoperable, and reusable) principles, all data are made accessible directly through the platform’s applications, both in their raw format and as processed subsets generated via the various online tools, together with any corresponding metadata. Additionally, supplemental files include key resources such as the ontology framework and other metadata used in building the platform, ensuring transparency and facilitating reuse by the community.

## Acknowledgments

The generation of the networks and all the bioinformatic analyses were performed on the HPC cluster Garnatxa at the Institute for Integrative Systems Biology (I2SysBio, UV-CSIC, Spain).

## Author contributions

A.S., L.O., D.N.-P., and J.T.M. participated in the design of the platform. A.Vi., I.M.-A., C.Z., P.L., D.P, C.M.-P., I.K., I.E., E.M., M.A.R.-S., A.Va., E.A.V. collected datasets relevant to their species of interest and research expertise. J.J. and A.D. developed the PETAL app. J.D.F. developed the GRN apps. A.S., L.O., and D.N.-P. developed the GCN apps and the remaining apps and platform structure. A.S., D. N.-P., and J.T.M. wrote the manuscript, and all authors contributed comments and suggestions.

## Supplemental Information

**Supplemental Figure 1.**
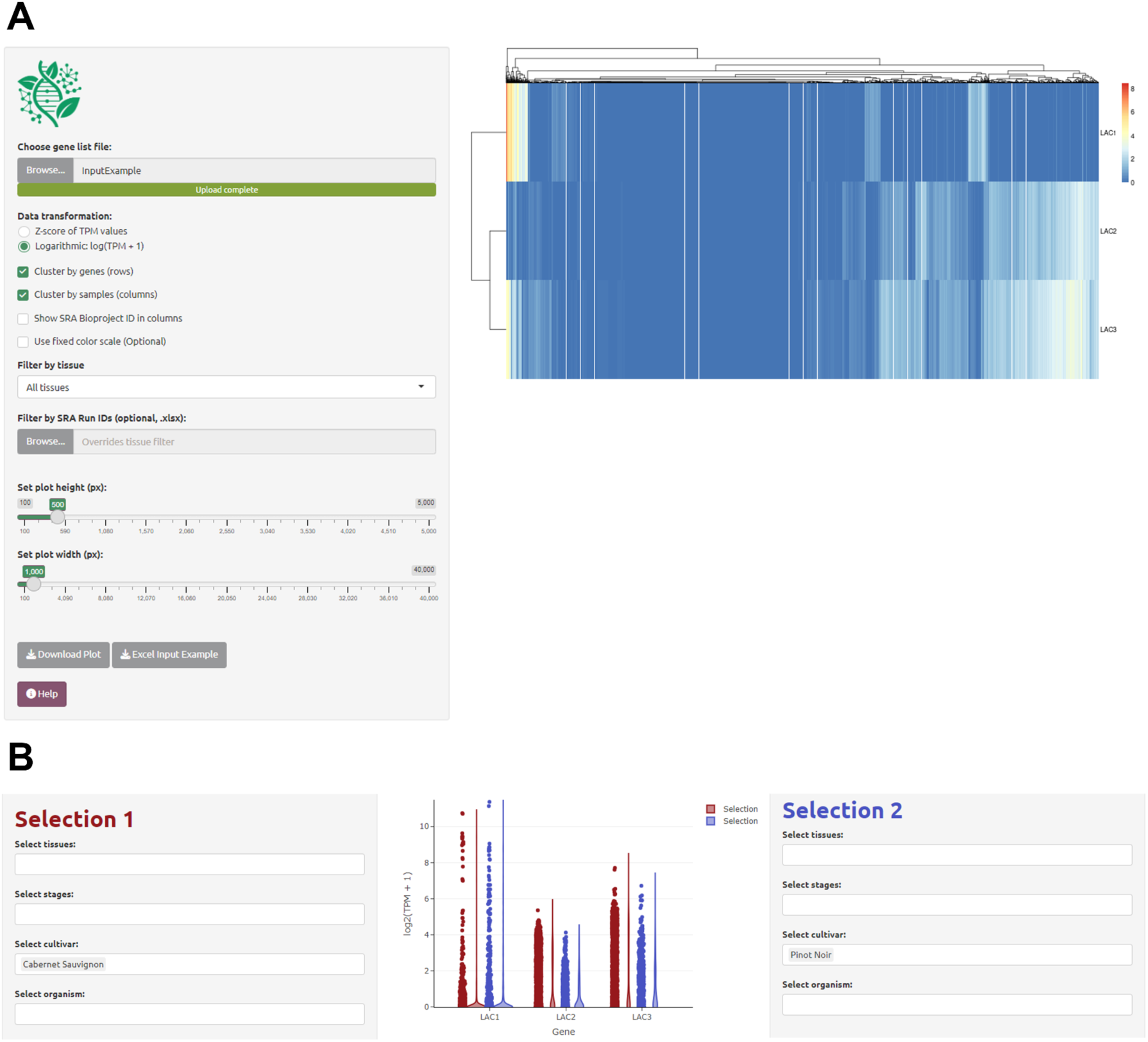
Example of *EXHARA* and *GeneXplore* expression comparison. (A) Expression heatmap across all SRA runs (*EXHARA*) for Vitis vinifera showing a different pattern for *vviLAC1*, *vviLAC2* and *vviLAC3*. (B) Differential expression profiles of *vviLAC1*, *vviLAC2*, and *vviLAC3* between two grapevine cultivars: cv. Cabernet Sauvignon (red) cv. Pinot Noir (blue). *GeneXplore* enables direct, customizable comparison of expression levels across selected genotypes, tissues, or any other category automatically classified via the platform’s ontologies. This feature facilitates the identification of transcriptional patterns and supports the formulation of functional hypotheses regarding gene roles that may only become apparent when looking at particular transcriptomic data subsets. Additional flexibility is provided by allowing filtering by metadata term confidence levels, as well as optionally expanding a query term to include all its child terms within the hierarchy.

**Supplemental Figure 2.**
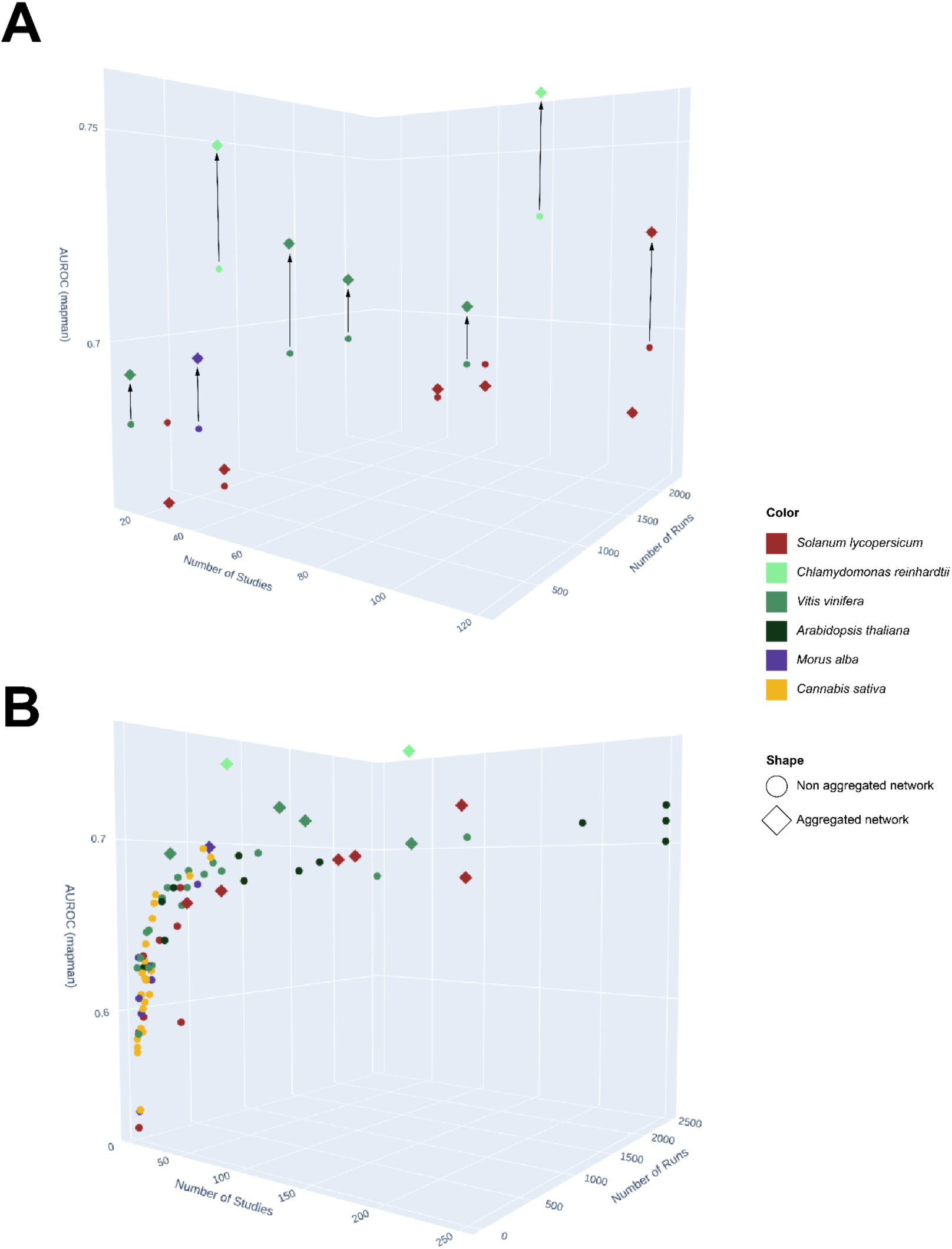
Impact of network aggregation and data volume on co-expression network performance. (A) Each vertical arrow connects a non-aggregated network (circle) to its corresponding aggregated network (diamonds) built from the same underlying data. Aggregated networks almost always outperform their non-aggregated counterparts across different species. Exceptions to this rule may be due to the selected tissue or that insufficient runs/studies were available. (B) All generated networks across a wide range of data volumes. This panel illustrates the general trend that network performance improves with an increasing number of studies and/or runs independent of the species within the platform or network method used. Note that lower AUROCs more often correspond to cannabis and mulberry networks, for which lower volumes of data are publicly available.

**Supplemental Figure 3.**
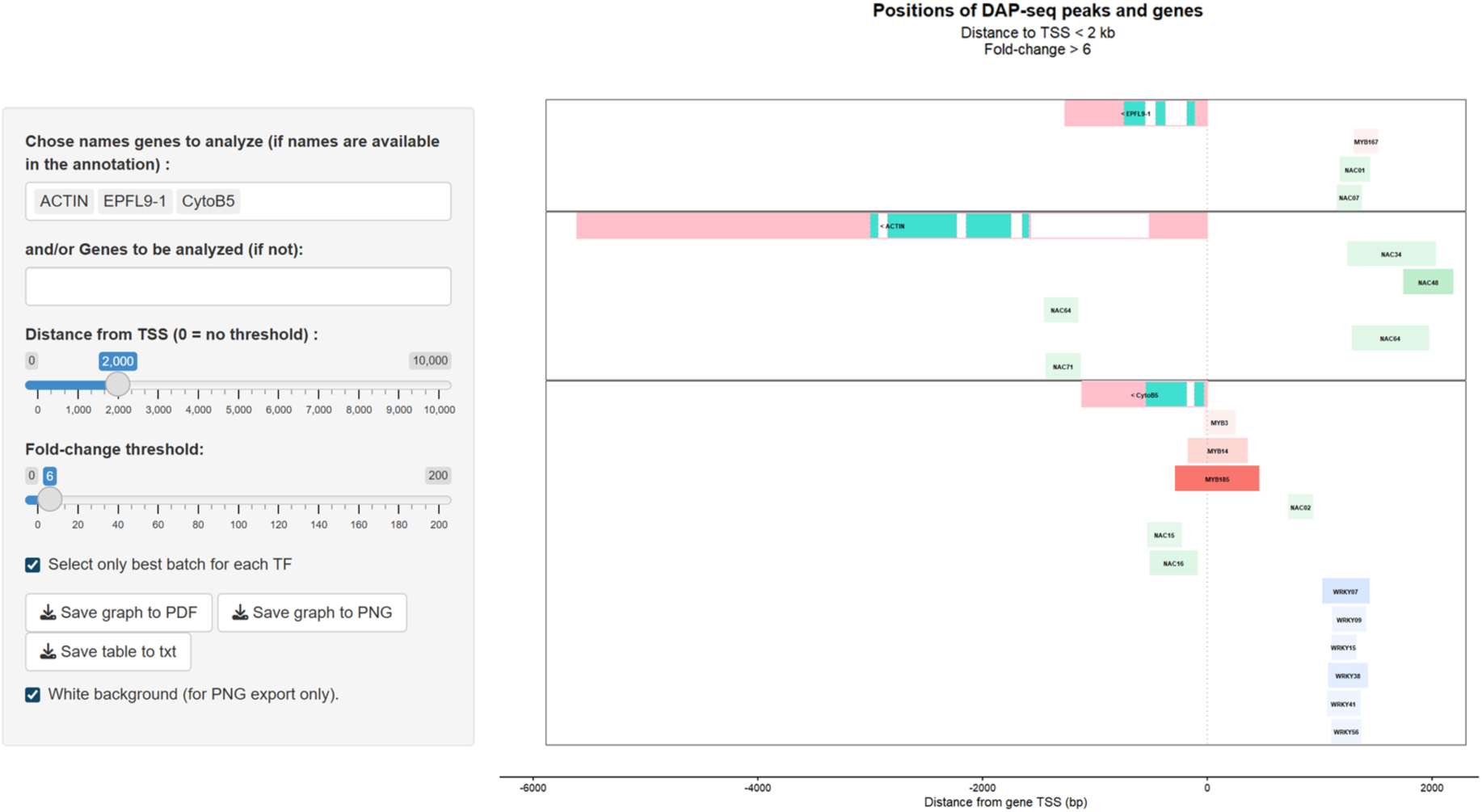
ChIP/DAP-seq peaks are visualized for three genes of interest (*vviACTIN*, *vviEPFL9-1*, and *vviCytoB5*) in the gene-centered tool of the *PETAL* app. These are displayed as individual gene plots centered on their respective transcriptional start site (TSS). Users can adjust threshold parameters for the maximum distance from the TSS (set here to 2 kb) and the minimum fold-change of peaks (set to >6). Only the highest-scoring peak batch for each TF is retained. Peaks are color-coded by TF-family, in this example, MYB (red), NAC (green), and WRKY (blue), with color intensity proportional to the fold-change score of the binding event. The promoter region analyzed for each gene is highlighted in pink. The application utilizes custom annotation files to render gene features and overlays peak information derived from DAP-seq data, all within an interactive Shiny interface. This enables intuitive exploration of TF–target gene interactions from a gene-centric view.

**Supplemental Figure 4.**
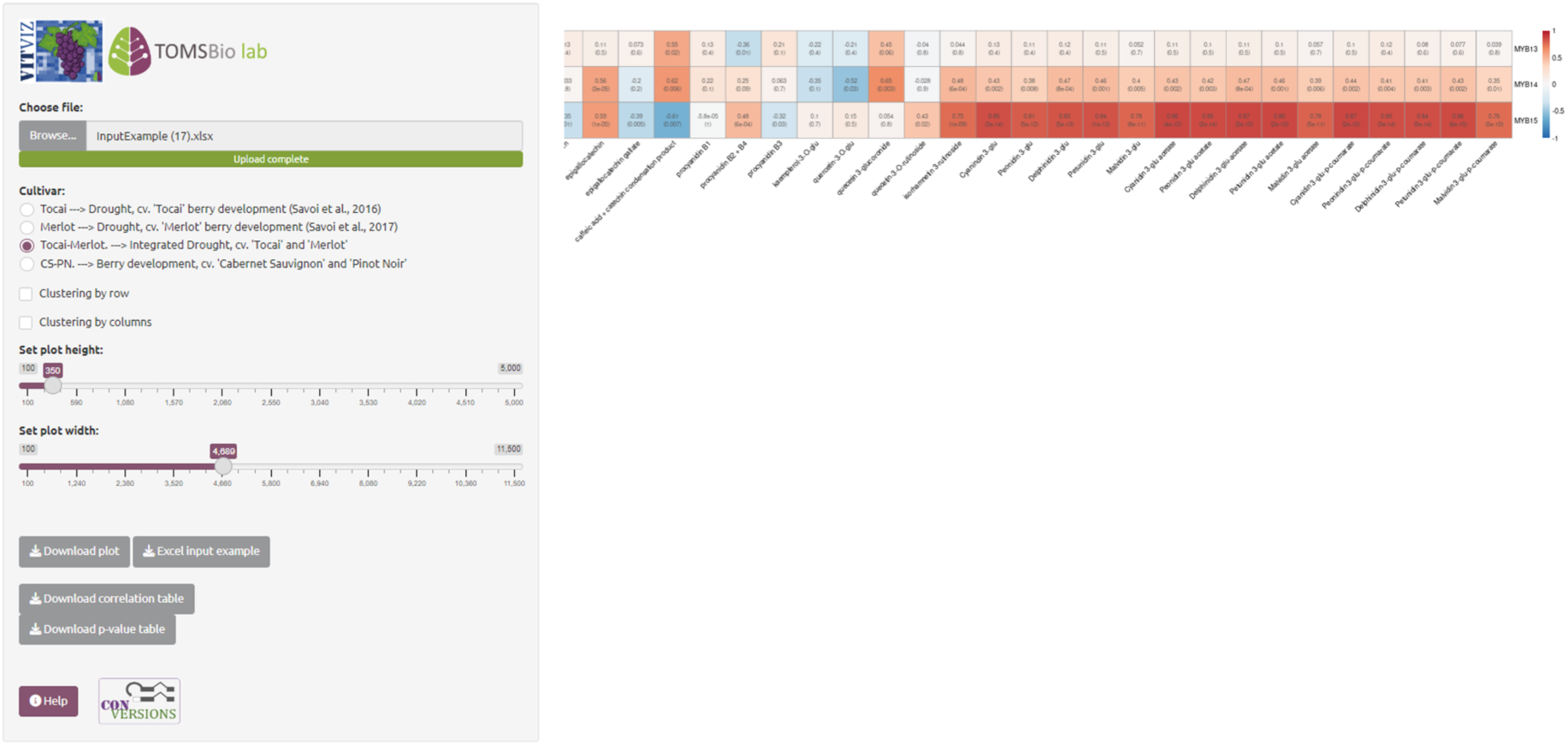
Example of *TransMetaDB* metabolite-transcript correlation analysis. Pearson correlation coefficients between transcript levels of *VviMYB13*, *VviMYB14*, and *VviMYB15* and multiple quantified metabolites in grape berry skins of *Vitis vinifera* cultivars ‘Tocai’ and ‘Merlot’ (Savoi et al., 2016, 2017). Each MYB gene shows distinct correlation profiles with subsets of metabolites, reflecting potential regulatory roles in cultivar-specific metabolic pathways. The app also allows for the use of a berry development dataset with cv. ‘Pinot Noir’ and ‘Cabernet Sauvignon’ samples (Fasoli et al., 2018). *TransMetaDB* enables such integrative analyses across paired transcriptomic and metabolomic datasets to support hypothesis generation on transcription factor function in metabolic regulation.

**Supplemental Figure 5.**
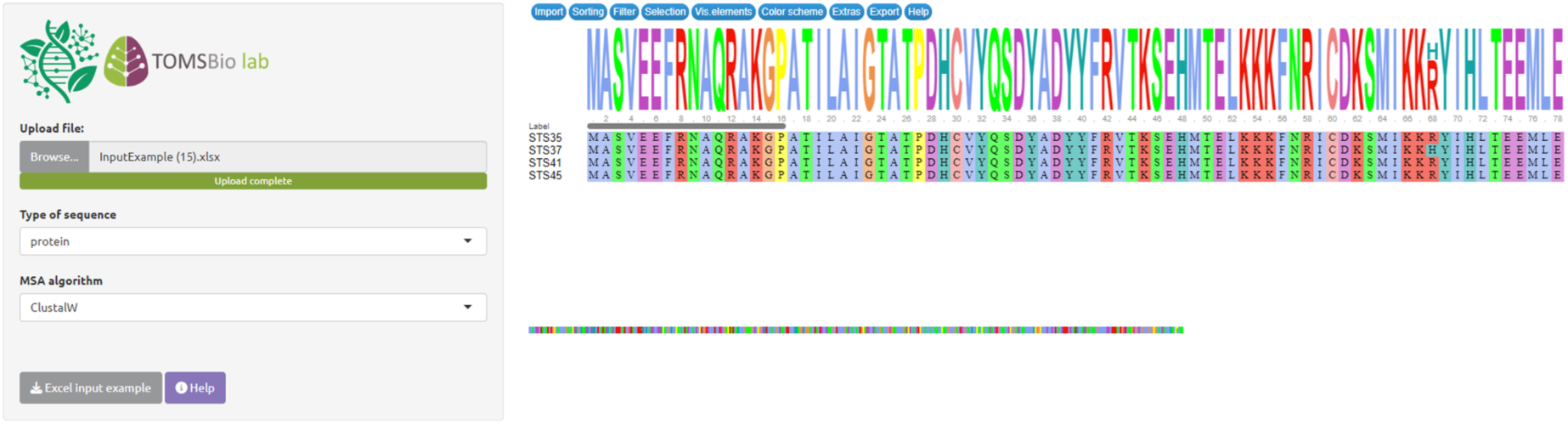
Example of MSA app. Alignment of VviSTS35, VviSTS37, VviSTS41 and VviSTS45 proteins.

**Supplemental Figure 6.**
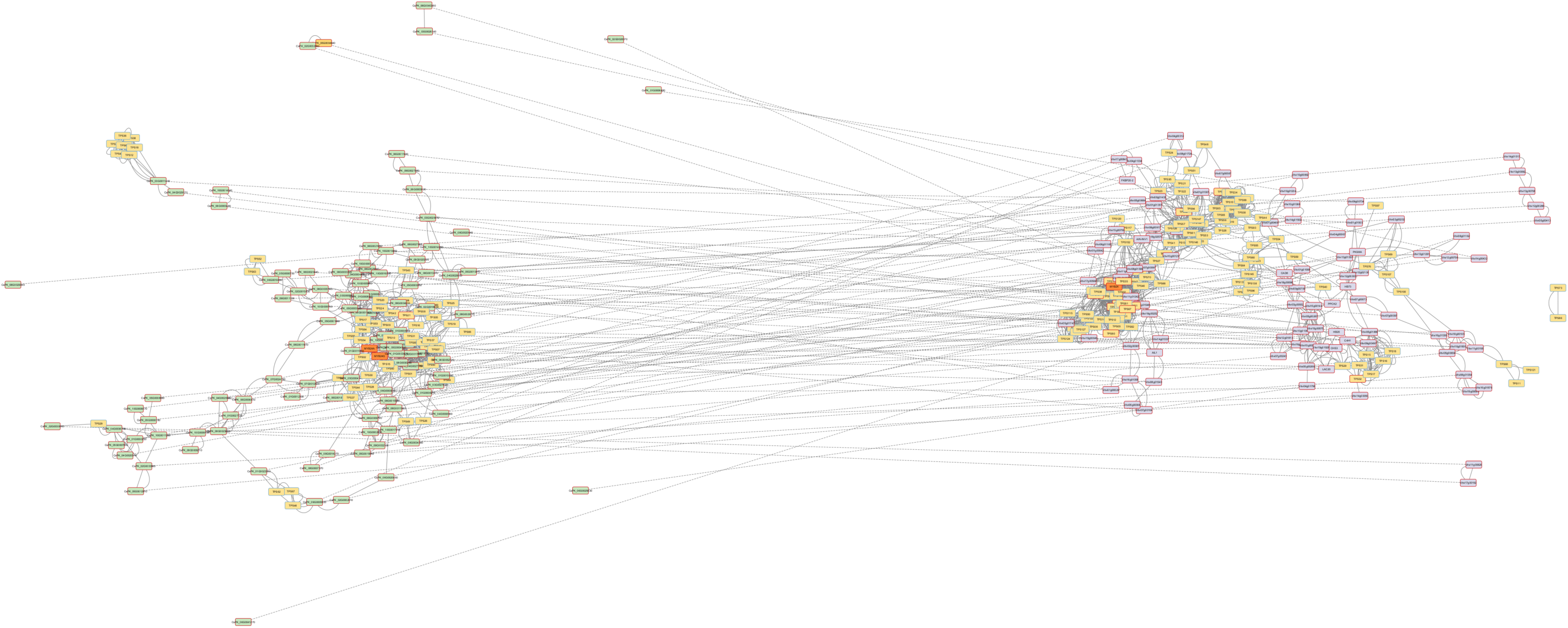
Coexpression networks exported from the GCN app and visualized in Cytoscape. (A) The network topology highlights both conserved and species-specific coexpression patterns. Overall topology similarities support a generally conserved role of MYB24 in terpene biosynthesis regulation in both species, even if some *MYB24-TPS* connections have changed from direct to indirect relationships, or *vice versa*. Node colors: green (C. sativa genes), purple (*V. vinifera* genes), yellow (*TPS* genes), and orange (*MYB24* genes). Red borders indicate DAP-seq binding. Dashed edges connect orthologous genes identified by reciprocal best BLAST hits (RBBH).

## References

Ballouz, S., Weber, M., Pavlidis, P., & Gillis, J. (2017). EGAD: Ultra-fast functional analysis of gene networks. Bioinformatics, 33(4), 612–614. 10.1093/bioinformatics/btw695

Carbonell-Bejerano, P., Rodríguez, V., Royo, C., Hernáiz, S., Moro-González, L., Torres-Viñals, M., & Martínez-Zapater, J. (2014). Circadian oscillatory transcriptional programs in grapevine ripening fruits. BMC Plant Biology, 14(1), 78. 10.1186/1471-2229-14-78

Chao, K.-H., Heinz, J. M., Hoh, C., Mao, A., Shumate, A., Pertea, M., & Salzberg, S. L. (2024). Combining DNA and protein alignments to improve genome annotation with LiftOn. Genome Research, genome;gr.279620.124v2. 10.1101/gr.279620.124

Childs, K. L., Davidson, R. M., & Buell, C. R. (2011). Gene Coexpression Network Analysis as a Source of Functional Annotation for Rice Genes. PLoS ONE, 6(7), e22196. 10.1371/journal.pone.0022196

Djari, A., Madignier, G., Di Valentin, O., Gillet, T., Frasse, P., Djouhri, A., Hu, G., Julliard, S., Liu, M., Zhang, Y., Regad, F., Pirrello, J., Maza, E., & Bouzayen, M. (2024). Haplotype-resolved genome assembly and implementation of VitExpress, an open interactive transcriptomic platform for grapevine. Proceedings of the National Academy of Sciences, 121(23). 10.1073/pnas.2403750121

Emms, D. M., & Kelly, S. (2019). OrthoFinder: Phylogenetic orthology inference for comparative genomics. Genome Biology, 20(1), 238. 10.1186/s13059-019-1832-y

Fasoli, M., Dal Santo, S., Zenoni, S., Tornielli, G. B., Farina, L., Zamboni, A., Porceddu, A., Venturini, L., Bicego, M., Murino, V., Ferrarini, A., Delledonne, M., & Pezzotti, M. (2012). The Grapevine Expression Atlas Reveals a Deep Transcriptome Shift Driving the Entire Plant into a Maturation Program. The Plant Cell, 24(9), 3489–3505. 10.1105/tpc.112.100230

Fasoli, M., Richter, C. L., Zenoni, S., Bertini, E., Vitulo, N., Dal Santo, S., Dokoozlian, N., Pezzotti, M., & Tornielli, G. B. (2018). Timing and Order of the Molecular Events Marking the Onset of Berry Ripening in Grapevine. Plant Physiology, 178(3), 1187–1206. 10.1104/pp.18.00559

Felemban, A., Moreno, J. C., Mi, J., Ali, S., Sham, A., AbuQamar, S. F., & Al-Babili, S. (2024). The apocarotenoid β-ionone regulates the transcriptome of *Arabidopsis thaliana* and increases its resistance against *Botrytis cinerea*. The Plant Journal, 117(2), 541–560. 10.1111/tpj.16510

Fernandez-Pozo, N., Menda, N., Edwards, J. D., Saha, S., Tecle, I. Y., Strickler, S. R., Bombarely, A., Fisher-York, T., Pujar, A., Foerster, H., Yan, A., & Mueller, L. A. (2015). The Sol Genomics Network (SGN)—From genotype to phenotype to breeding. Nucleic Acids Research, 43(D1), D1036–D1041. 10.1093/nar/gku1195

Goodstein, D. M., Shu, S., Howson, R., Neupane, R., Hayes, R. D., Fazo, J., Mitros, T., Dirks, W., Hellsten, U., Putnam, N., & Rokhsar, D. S. (2012). Phytozome: A comparative platform for green plant genomics. Nucleic Acids Research, 40(D1), D1178–D1186. 10.1093/nar/gkr944

Guo, Y., Mahony, S., & Gifford, D. K. (2012). High Resolution Genome Wide Binding Event Finding and Motif Discovery Reveals Transcription Factor Spatial Binding Constraints. PLoS Computational Biology, 8(8), e1002638. 10.1371/journal.pcbi.1002638

Huala, E., Dickerman, A. D., Garcia-Hernandez, M., Weems, D., Reiser, L., LaFond, F., Hanley, D., Kiphart, D., Zhuang, M., Huang, W., Mueller, L. A., Bhattacharyya, D., Bhaya, D., Sobral, B. W., Beavis, W., Meinke, D. W., Town, C. D., Somerville, C., & Yon Rhee, S. (2001). The Arabidopsis Information Resource (TAIR): A comprehensive database and web-based information retrieval, analysis, and visualization system for a model plant. Nucleic Acids Research, 29(1), 102–105. 10.1093/nar/29.1.102

Huynh-Thu, V. A., Irrthum, A., Wehenkel, L., & Geurts, P. (2010). Inferring Regulatory Networks from Expression Data Using Tree-Based Methods. PLoS ONE, 5(9), Article 9. 10.1371/journal.pone.0012776

Langmead, B., & Salzberg, S. L. (2012). Fast gapped-read alignment with Bowtie 2. Nature Methods, 9(4), 357–359. 10.1038/nmeth.1923

Li, H. (2023). Protein-to-genome alignment with miniprot. Bioinformatics, 39(1), btad014. 10.1093/bioinformatics/btad014

Liesecke, F., De Craene, J.-O., Besseau, S., Courdavault, V., Clastre, M., Vergès, V., Papon, N., Giglioli-Guivarc’h, N., Glévarec, G., Pichon, O., & Dugé De Bernonville, T. (2019). Improved gene co-expression network quality through expression dataset down-sampling and network aggregation. Scientific Reports, 9(1). 10.1038/s41598-019-50885-8

Liu, S., Zhong, Z., Sun, Z., Tian, J., Sulaiman, K., Shawky, E., Fu, H., & Zhu, W. (2022). *De novo* Transcriptome Analysis Revealed the Putative Pathway Genes Involved in Biosynthesis of Moracins in *Morus alba* L. ACS Omega, 7(13), 11343–11352. 10.1021/acsomega.2c00409

Livingston, S. J., Quilichini, T. D., Booth, J. K., Wong, D. C. J., Rensing, K. H., Laflamme-Yonkman, J., Castellarin, S. D., Bohlmann, J., Page, J. E., & Samuels, A. L. (2020). Cannabis glandular trichomes alter morphology and metabolite content during flower maturation. The Plant Journal, 101(1), 37–56. 10.1111/tpj.14516

Navarro-Payá, D., Santiago, A., Orduña, L., Zhang, C., Amato, A., D’Inca, E., Fattorini, C., Pezzotti, M., Tornielli, G. B., Zenoni, S., Rustenholz, C., & Matus, J. T. (2022). The Grape Gene Reference Catalogue as a Standard Resource for Gene Selection and Genetic Improvement. Frontiers in Plant Science, 12, 803977. 10.3389/fpls.2021.803977

Orduña, L., Santiago, A., Navarro-Payá, D., Zhang, C., Wong, D. C. J., & Matus, J. T. (2023). Aggregated gene co-expression networks predict transcription factor regulatory landscapes in grapevine. Journal of Experimental Botany, 74(21), Article 21. 10.1093/jxb/erad344

Priyam, A., Woodcroft, B. J., Rai, V., Moghul, I., Munagala, A., Ter, F., Chowdhary, H., Pieniak, I., Maynard, L. J., Gibbins, M. A., Moon, H., Davis-Richardson, A., Uludag, M., Watson-Haigh, N. S., Challis, R., Nakamura, H., Favreau, E., Gómez, E. A., Pluskal, T., … Wurm, Y. (2019). Sequenceserver: A Modern Graphical User Interface for Custom BLAST Databases. Molecular Biology and Evolution, 36(12), 2922–2924. 10.1093/molbev/msz185

Reiser, L., Bakker, E., Subramaniam, S., Chen, X., Sawant, S., Khosa, K., Prithvi, T., & Berardini, T. Z. (2024). The Arabidopsis Information Resource in 2024. Genetics, 227(1), iyae027. doi10.1093

Santiago, A., Romero, P., Martínez, A., Martínez, M. J., Echeverría, J., Selles, S., Alvarez-Urdiola, R., Zhang, C., Navarro-Payá, D., Pizzio, G. A., Micó, E., Samper, A., Morante, J., Cigliano, R. A., Manzano, D., Bru, R., & Matus, J. T. (2024). *Biosynthesis of oxyresveratrol in mulberry (* Morus alba *L.) is mediated by a group of p-coumaroyl-CoA 2’-hydroxylases acting upstream of stilbene synthases*. 10.1101/2024.04.04.588114

Savoi, S., Santiago, A., Orduña, L., & Matus, J. T. (2022). Transcriptomic and metabolomic integration as a resource in grapevine to study fruit metabolite quality traits. Frontiers in Plant Science, 13, 937927. 10.3389/fpls.2022.937927

Savoi, S., Wong, D. C. J., Arapitsas, P., Miculan, M., Bucchetti, B., Peterlunger, E., Fait, A., Mattivi, F., & Castellarin, S. D. (2016). Transcriptome and metabolite profiling reveals that prolonged drought modulates the phenylpropanoid and terpenoid pathway in white grapes (Vitis vinifera L.). BMC Plant Biology, 16(1). 10.1186/s12870-016-0760-1

Savoi, S., Wong, D. C. J., Degu, A., Herrera, J. C., Bucchetti, B., Peterlunger, E., Fait, A., Mattivi, F., & Castellarin, S. D. (2017). Multi-Omics and Integrated Network Analyses Reveal New Insights into the Systems Relationships between Metabolites, Structural Genes, and Transcriptional Regulators in Developing Grape Berries (Vitis vinifera L.) Exposed to Water Deficit. Frontiers in Plant Science, 8. 10.3389/fpls.2017.01124

Sayers, E. W., Bolton, E. E., Brister, J. R., Canese, K., Chan, J., Comeau, D. C., Connor, R., Funk, K., Kelly, C., Kim, S., Madej, T., Marchler-Bauer, A., Lanczycki, C., Lathrop, S., Lu, Z., Thibaud-Nissen, F., Murphy, T., Phan, L., Skripchenko, Y., … Sherry, S. T. (2022). Database resources of the national center for biotechnology information. Nucleic Acids Research, 50(D1), D20– D26. 10.1093/nar/gkab1112

Shumate, A., & Salzberg, S. L. (2021). Liftoff: Accurate mapping of gene annotations. Bioinformatics, 37(12), 1639–1643. 10.1093/bioinformatics/btaa1016

Szklarczyk, D., Kirsch, R., Koutrouli, M., Nastou, K., Mehryary, F., Hachilif, R., Gable, A. L., Fang, T., Doncheva, N. T., Pyysalo, S., Bork, P., Jensen, L. J., & von Mering, C. (2023). The STRING database in 2023: Protein–protein association networks and functional enrichment analyses for any sequenced genome of interest. Nucleic Acids Research, 51(D1), D638–D646. 10.1093/nar/gkac1000

Vannozzi, A., Palumbo, F., Magon, G., Lucchin, M., & Barcaccia, G. (2021). The grapevine (Vitis vinifera L.) floral transcriptome in Pinot noir variety: Identification of tissue-related gene networks and whorl-specific markers in pre- and post-anthesis phases. Horticulture Research, 8(1). 10.1038/s41438-021-00635-7

Wong, D. C. J. (2020). Network aggregation improves gene function prediction of grapevine gene co-expression networks. Plant Molecular Biology, 103(4–5), 425–441. 10.1007/s11103-020-01001-2

Zhang, C., Dai, Z., Ferrier, T., Orduña, L., Santiago, A., Peris, A., Wong, D. C. J., Kappel, C., Savoi, S., Loyola, R., Amato, A., Kozak, B., Li, M., Liang, A., Carrasco, D., Meyer-Regueiro, C., Espinoza, C., Hilbert, G., Figueroa-Balderas, R., … Matus, J. T. (2023). MYB24 orchestrates terpene and flavonol metabolism as light responses to anthocyanin depletion in variegated grape berries. The Plant Cell, 35(12), 4238–4265. 10.1093/plcell/koad228

Zhu, L. J., Gazin, C., Lawson, N. D., Pagès, H., Lin, S. M., Lapointe, D. S., & Green, M. R. (2010). ChIPpeakAnno: A Bioconductor package to annotate ChIP-seq and ChIP-chip data. BMC Bioinformatics, 11(1), 237. 10.1186/1471-2105-11-237

Zouine, M., Maza, E., Djari, A., Lauvernier, M., Frasse, P., Smouni, A., Pirrello, J., & Bouzayen, M. (2017). TomExpress, a unified tomato RNA-Seq platform for visualization of expression data, clustering and correlation networks. The Plant Journal, 92(4), 727–735. 10.1111/tpj.13711

