## Supplementary figures and images for "PlantaeViz: a comprehensive platform for the analysis, integration, and visualization of public omics data across plant and related species"

### Supplemental Data 6.pdf

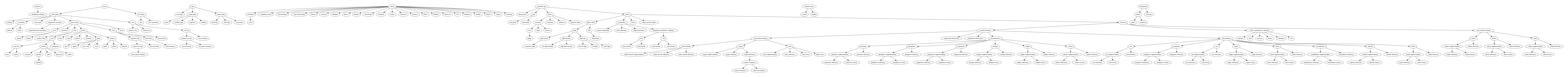

### Supplemental Data 7.pdf

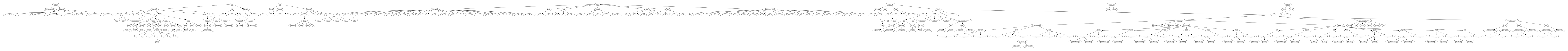

### Supplemental Data 9.pdf

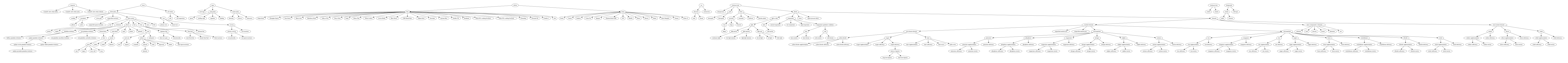
